# Reward-based improvements in motor control are driven by multiple error-reducing mechanisms

**DOI:** 10.1101/792598

**Authors:** Olivier Codol, Peter J. Holland, Sanjay G. Manohar, Joseph M. Galea

## Abstract

Reward has a remarkable ability to invigorate motor behaviour, enabling individuals to select and execute actions with greater precision and speed. However, if reward is to be exploited in applied settings such as rehabilitation, a thorough understanding of its underlying mechanisms is required. Although reward-driven enhancement of movement execution has been proposed to occur through enhanced feedback control, an untested alternative is that it is driven by increased arm stiffness, an energy-consuming process that increases limb stability. First, we demonstrate that during reaching reward improves selection and execution performance concomitantly without interference. Computational analysis revealed that reward led to both an increase in feedback correction during movement and a reduction in motor noise near the target. We provide novel evidence that this noise reduction is driven by a reward-dependent increase in arm stiffness. Therefore, reward drives multiple error-reduction mechanisms which enable individuals to invigorate motor performance without compromising accuracy.

## 1 Introduction

Motor control involves two main components that may be individually optimised, action selection and action execution (Chen, Holland & Galea, 2018). While the former addresses the problem of finding the best action to achieve a goal amongst a subset of actions, the latter is concerned with performing the selected action with the greatest precision possible (Chen, Holland & Galea, 2018; Shmuelof et al., 2014; Stanley & Krakauer, 2013). Naturally, both processes come at a computational cost, meaning the faster an action is selected or executed, the more prone it is to errors – a phenomenon formalised as Fitts’ law (Fitts, 1954). This is represented in a speed-accuracy function where accuracy decays as speed increases. Because speed-accuracy functions are a hallmark of human limitation in motor control, they have been regularly used to quantify performance (Reis et al., 2009; Telgen et al., 2014). For example, in skill learning, one may see the speed-accuracy function shift so that higher levels of accuracy are observed for any given speed (Reis et al., 2009; Telgen et al., 2014).

Interestingly, both action selection and action execution are highly susceptible to the presence of reward. For instance, introducing monetary reward in a sequence learning task leads to a reduction in selection errors, as well as a decrease in reaction times, suggesting faster computation at no cost to accuracy (Wachter et al., 2009). Similarly, in a saccade task, reward reduced participant’s reaction time whilst making them less sensitive to distractors (Manohar et al., 2015). It has also been shown that reward invigorates movement execution by increasing peak velocity and accuracy during saccades (Manohar et al., 2015; Takikawa et al., 2002) and reaching movements (Carroll et al., 2019; Galaro et al., 2019; Summerside et al., 2018). Therefore, this body of work suggests that reward can consistently shift the speed-accuracy function, at least in isolation, of both selection and execution. It has also been shown that in saccades reward can enhance the selection and execution components concomitantly (Manohar et al., 2015). However, it is currently unclear whether this generalizes to more complex reaching movements. As the use of reward has generated much interest as a potential tool to enhance rehabilitation procedures for clinical populations (Goodman et al., 2014; Quattrocchi et al., 2017), it is crucial to determine whether reward can improve the selection and execution components of a reaching movement without interference.

Another open question is how reward mechanistically drives improvements in performance. Recent work in eye and reaching movements suggests that reward acts by increasing feedback control, enhancing one’s ability to correct for movement error (Carroll et al., 2019; Manohar et al., 2019). However, there are far simpler mechanisms which reward could utilize to improve execution. For example, the motor system has the ability to control the stiffness of its effectors, such as the arm during a reaching task, by employing co-contraction of ant-agonist muscles at once (Gribble et al., 2003; Perreault et al., 2002). This increase in arm stiffness results in the limb being more stable in the face of perturbations (Franklin et al., 2007), and capable of absorbing noise that may arise during the movement itself (Selen et al., 2009; Ueyama & Miyashita, 2013), thus reducing error and improving performance (Gribble et al., 2003). Yet, it is unclear whether the reward-based improvements in execution are related to increased arm stiffness.

To address these questions, we devised a reaching task in which participants could be rewarded with money as a function of their reaction time and movement time. Occasionally, distractor targets of a different colour appeared, and participants were told to withhold movement until the correct target subsequently appeared, allowing for a selection component to be quantified. In a first experiment, we show that reward improves both selection and execution concomitantly, and that the presence or absence of reward, rather than reward magnitude modulated this effect. In a second experiment, we asked whether punishment had a similar effect to reward. We demonstrate that although both reward and punishment led to similar effects, action execution, but not action selection, showed a more global, non-contingent sensitivity to punishment. Behavioural and computational analysis suggested that in addition to an increase in feedback corrections during movement, reward may have improved motor execution through an increase in arm stiffness leading to a decrease in motor noise at the end of the movement. In a third and fourth experiment, we tested this hypothesis and provide evidence that this reduction in noise is driven by a reward-dependent increase in arm stiffness. Therefore, reward not only invigorates motor execution performance by increasing the contribution of feedback control, but also protects against noise at the peripheral level via an increase in arm stiffness.

## 2 Results

### 2.1 Reward concomitantly enhances action selection and action execution

Experiment 1 examined the effect of reward on the selection and execution components of a reaching movement. Whilst holding a robotic manipulandum, participants (N=30) made discrete reaching movements towards 1 of 4 visual targets presented 20cm away from a central start position (figure 1A). To assess the effect of reward value on reaching performance, participants were informed of the upcoming trial type prior to movement onset: 0p, 10p and 50p. For the 10p and 50p trials participants could earn money based on their combined reaction time and movement time. The scoring function which translated performance to monetary gain was adaptive (figure 1B), factoring in the recent history of movement times and reaction times to ensure participants experienced comparable amounts of reward despite idiosyncrasies in individual’s reaction times and movement speed (Berret et al., 2018; Reppert et al., 2018; Manohar et al., 2015). To assess selection and execution performance concomitantly, we interleaved normal trials and distractor trials. In normal trials, the target’s colour matched the starting position colour (figure 1C), while in distractor trials (42% of trials) a distractor target bearing a different colour than the starting position appeared prior to the correct target (figure 1D). In this case, participants were instructed to withhold their movement to the distractor and wait until the correct target appeared before making a movement. If participants exited the starting position upon appearance of a distractor, the trial was considered as “distracted”. While the probability of initiating reaches to a distractor target provided a measure of selection accuracy, the associated reaction times provided a selection speed, allowing us to define a speed-accuracy function (Fitts, 1954; Hübner & Schlösser, 2010; Manohar et al., 2015). For execution, radial error provided a measure of execution accuracy while peak velocity during the reach and movement time provided an execution speed, again allowing us to define a speed-accuracy function.

**Figure 1.**
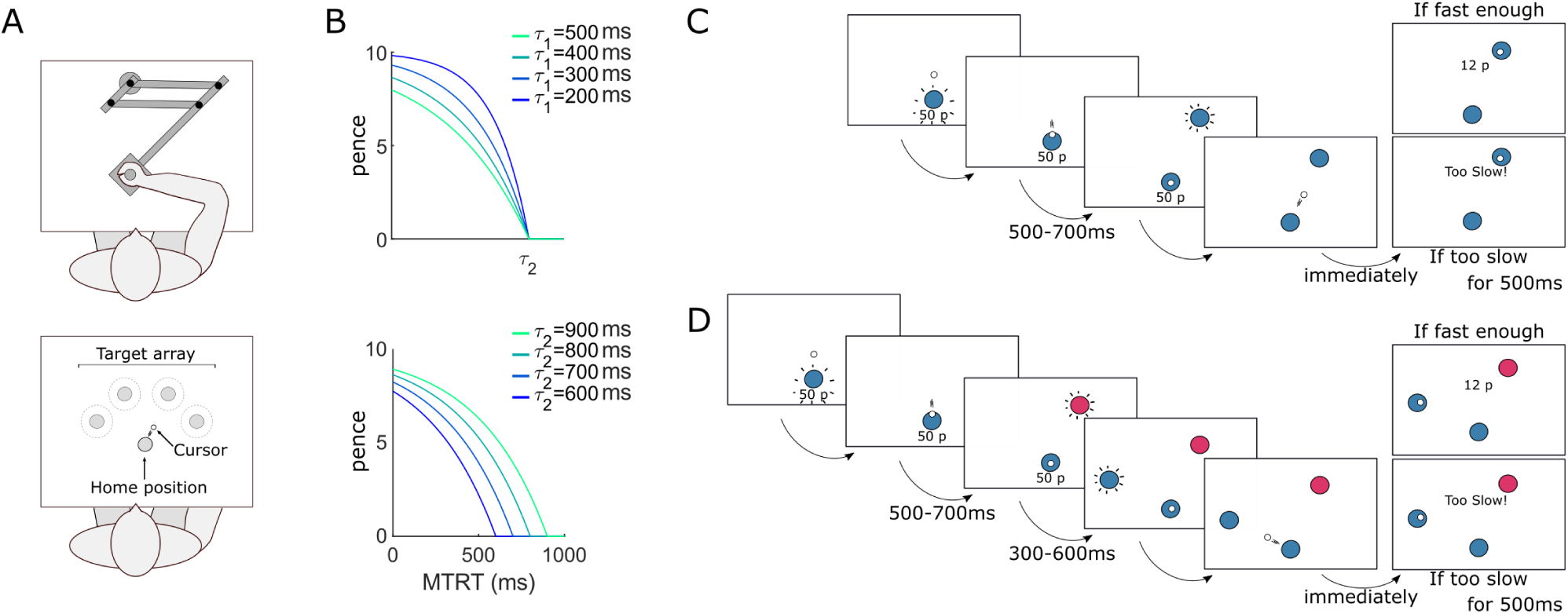
Reaching paradigm. A. Participants reached to a series of targets using a robotic manipulandum. B. The faster participants moved, the more money they made. Speed was the sum of movement time and reaction time (MTRT) and the function varied based on two parameters *τ*_1_ and *τ*_2_. The upper and lower plots show how the function varied as a function of *τ*_1_ (*τ*_2_ fixed at 800ms) and *τ*_2_ (*τ*_1_ fixed at 400ms), respectively, for a 10p trial. C. Normal trial. Participants reached at a single target and earned money based on their performance speed. If they were too slow (MTRT*<τ*_2_), a message “*Too slow!* “ appeared instead of the reward information. Transition times are indicated below for each screen. A uniform distribution was employed for the transition time jitter. D. Distractor trial. Occasionally, a first target bearing a different colour appeared, and participants were told to wait for the second, correct target to appear and reach toward the latter.

To analyse if speed-accuracy functions were altered by reward, trials for each reward value and participant were sorted as a function of their speed (reaction time for selection and peak velocity for execution) and divided into 50 quantiles (Manohar et al., 2015). For each quantile, the average accuracy (percentage of non-distracted trials and radial error) over a 30% centile window was obtained. Group averages were then obtained for each quantile in the speed and accuracy dimension, and results are displayed in figure 2. As expected, reward shifted the speed-accuracy functions for both selection and execution, underlining augmented motor performance with reward.

**Figure 2.**
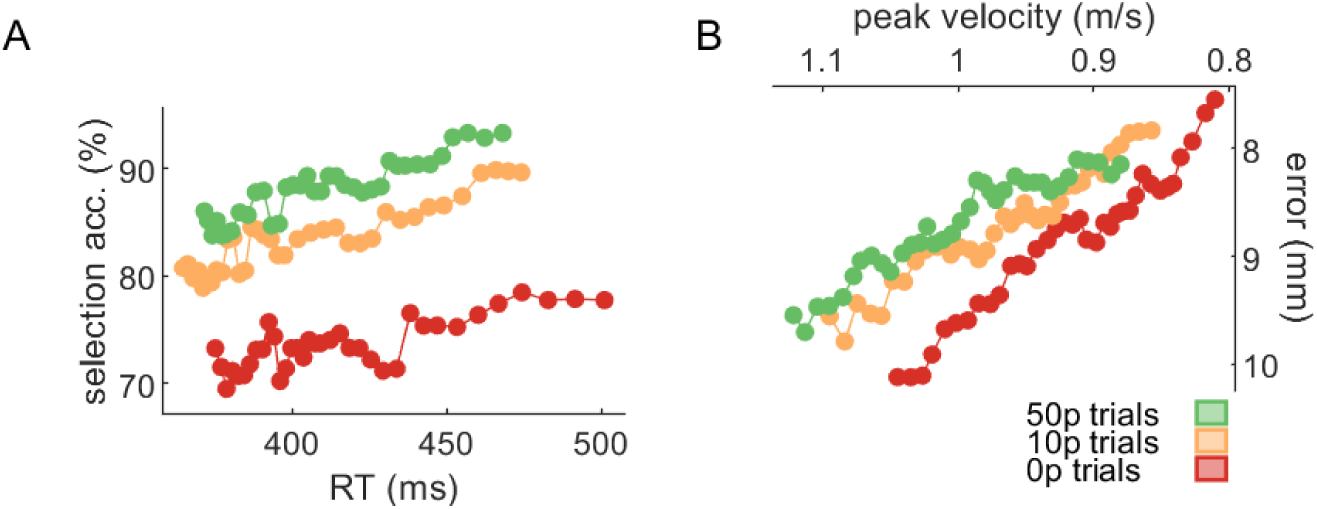
Speed-accuracy functions for selection (A) and execution (B) shift as reward values increase. The functions are obtained by sliding a 30% centile window over 50 quantile-based bins. A. For the selection panel, the count of non-distracted trials and distracted trials for each bin was obtained, and the ratio (100*non-distracted/total) calculated afterwards. B. For the execution component, the axes were inverted to match the selection panel in A, *i.e.* the upper left corner indicates faster and more accurate performance. See methods section Dataanalysis and text for details.

Comparing each variable of interest individually, participants showed a clear and consistent improvement in selection accuracy in the presence of reward. Specifically, they were less likely to be distracted in rewarded trials, though this was independent of reward magnitude (repeated-measures ANOVA, *F* (2) = 15.8, *p <* 0.001, partial *η*^2^ = 0.35, *post-hoc* 0p vs 10p *t*(29) = −3.34, *p* = 0.005, *d* = −0.61; 0p vs 50p *t*(29) = −5.32, *p <* 0.001, *d* = −0.97; 10p vs 50p *t*(29) = −2.21, *p* = 0.07, *d* = −0.49; figure 3A). However, this did not come at the cost of slowed decision-making, as reaction times remained largely similar across reward values; if anything, reaction times were slightly shorter if a large reward (50p) was available compared to no-reward (0p) trials, though this was not statistically significant (*F* (2) = 2.35, *p* = 0.10, partial *η*^2^ = 0.07; figure 3B-C).

**Figure 3.**
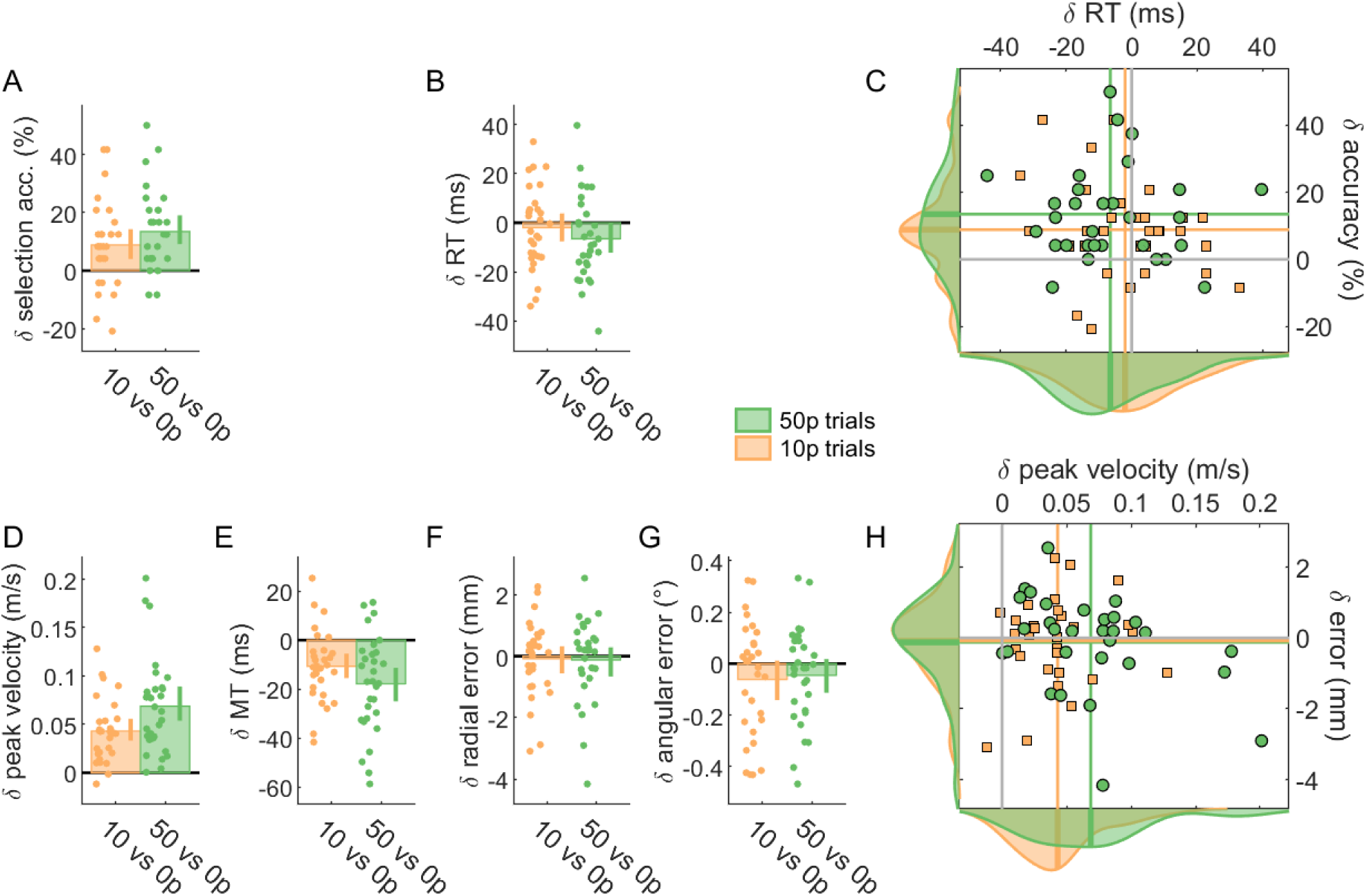
Reward enhances performance in both selection and execution. For all bar plots, data was normalised to 0p performance for each individual. Bar height indicates group mean, dots represent individual values and error bars indicate bootstrapped 95% CIs of the mean. A. Selection accuracy, as the percentage of trials where participants initiated reaches toward the correct target instead of the distractor target. B. Mean reaction times. C. Scatterplot of mean reaction time against selection accuracy. Values are normalised to 0p trials. The coloured lines indicate the mean value for each condition, and the solid grey lines indicate the origin, that is, 0p performance. Data distributions are displayed on the sides, with transversal bars indicating the mean of the distribution. Triangles indicate 50p trials. D. Mean peak velocity during reaches. E. Mean movement times of reaches. F. Mean radial error at the end of the reach. G. Mean angular error at the end of the reach. H. Scatterplot showing execution speed (peak velocity) against execution accuracy (radial error), similar to C.

In addition, reward led to a marked improvement in action execution by increasing peak velocity that scaled with reward magnitude, although this was driven by three extreme values (*F* (2) = 43.0, *p <* 0.001, partial *η*^2^ = 0.60, *post-hoc* 0p vs 10p *t*(29) = −7.40, *p <* 0.001, *d* = −1.35; 0p vs 50p *t*(29) = −7.61, *p <* 0.001, *d* = −1.39; 10p vs 50p *t*(29) = −3.52, *p* = 0.003, *d* = −0.64; figure 3D). Unsurprisingly, movement time also showed a similar effect, that is, mean movement time decreased with reward, though this did not scale with reward magnitude (*F* (2) = 15.3, *p <* 0.001, partial *η*^2^ = 0.35, *post-hoc* 0p vs 10p *t*(29) = 4.07, *p <* 0.001, *d* = 0.74; 0p vs 50p *t*(29) = 4.99, *p <* 0.001, *d* = 0.91; 10p vs 50p *t*(29) = 2.08, *p* = 0.09, *d* = 0.38; figure 3E). However, this reward-based improvement in speed did not come at the cost of accuracy as radial error (*F* (2) = 0.15, *p* = 0.86, partial *η*^2^ = 0.005) and angular error (*F* (2) = 1.51, *p* = 0.23, partial *η*^2^ = 0.05) remained unchanged (figure 3F-H).

These results demonstrate that reward enhanced the selection and execution components of a reaching movement simultaneously and without interference. Interestingly, these improvements were mainly driven by an increase in accuracy for selection and in speed for execution. However, reward magnitude had only a marginal impact on the effect of reward itself, as opposed to the presence or absence of reward *per se*. Consequently, for the remaining studies, we used the 0p and 50p trial conditions to assess the impact of reward on reaching performance.

### 2.2 Punishment has the same effect as reward on selection but a non-contingent effect on execution

Next, we asked if punishment led to the same effect as reward, as previous reports have shown that they have dissociable effects on motor performance (Galea et al., 2015; Hamel et al., 2018; Song & Smiley-Oyen, 2017; Wachter et al., 2009). A new group of participants (N=30) experienced a reward and a punishment block in a counterbalanced order. In the reward block, 0p and 50p trials were randomly interleaved. Similar to the previous experiment, on 50p trials participants received money as a result of fast reaction times and movement times. The punishment block consisted of randomly interleaved -0p and -50p trials which indicated the maximum amount of money that could be lost on a single trial. At the beginning of this block, participants were given £11, and on -50p trials, participants lost money as a result of slow reaction times and movement times.

First, we obtained speed-accuracy functions for the selection and execution components in the same way as for experiment 1 (figure 4). While punishment had a similar effect on selection (Figure 4A), it produced dissociable effects on execution (Figure 4B). Specifically, while peak velocity increased with punishment similarly to reward, it was accompanied by an increase in radial error. Although this could suggest that punishment does not cause a change in the speed-accuracy function relative to its own baseline (−0p) trials, a clear shift in the speed-accuracy function could be seen between the baseline trials of the reward and punishment conditions (Figure 4B). Therefore, relative to reward, a punishment context appeared to have a non-contingent beneficial effect on motor execution.

**Figure 4.**
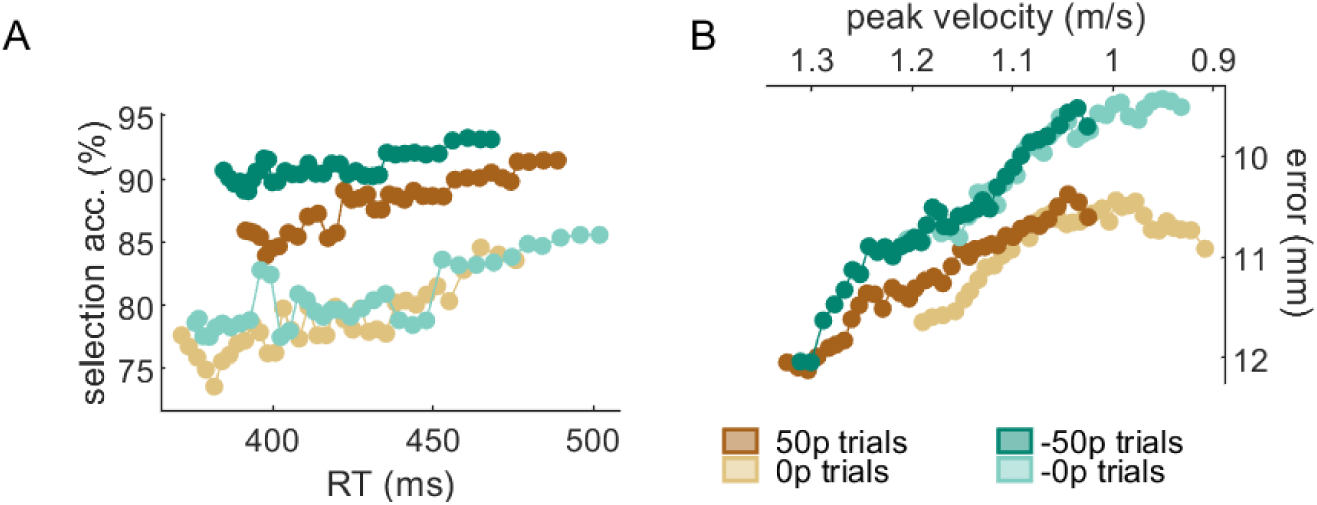
Reward and punishment speed-accuracy functions for selection (A) and execution (B) components. The functions are obtained by sliding a 30% centile window over 50 quantile-based bins. A. For the selection panel, the count of non-distracted trials and distracted trials for each bin was obtained, and the ratio (100*non-distracted/total) calculated afterwards. B. For the execution component, the axes were inverted to match the selection panel in A, *i.e.* the upper left corner indicate faster and more accurate performance. See methods section Data analysis and text for details.

To examine these results further, we fitted a mixed-effect linear model *DV* ∼ 1 + *RP* + *value* + *RP* : *value* + (1|*participant*) that included individual intercepts and an interaction term, where *DV* is the dependent variable considered, *RP* indicated whether the context was reward or punishment (*i.e.* reward block or punishment block) and *value* indicated whether the trial is a baseline trial bearing no value (0p and -0p) or a rewarded/punished trial bearing high value (50p and -50p). as in experiment 1, *value* improved selection accuracy (*β* = 9.72, CI = [4.51, 14.9], *t*(116) = 3.70, *p <* 0.001; figure 5A) without any effect on reaction times (*β* = −0.007, CI = [−0.015, 0.002], *t*(116) = −1.53, *p* = 0.13; figure 5B,C) and increased peak velocity and decreased movement time (main effect of value on peak velocity *β* = 0.096, CI = [0.045, 0.147], *t*(116) = 3.76, *p <* 0.001; on movement time *β* = −0.02, CI = [−0.033, −0.007], *t*(116) = −3.15, *p* = 0.002; figure 5D,E) at no accuracy cost (radial error *β* = −0.085, CI = [−0.001, 0.171], *t*(116) = 1.96, *p* = 0.052; angular error *β* = 0.081, CI = [−0.027, 0.189], *t*(116) = 1.49, *p* = 0.14; figure 5F-H), therefore replicating the findings from experiment 1. Importantly, context (reward vs. punishment) did not alter these effects on selection accuracy (main effect of block *β* = −1.94, CI = [−7.15, 3.26], *t*(116) = −0.74, *p* = 0.46; interaction *β* = −0.97, CI = [−8.34, 6.39], *t*(116) = −0.26, *p* = 0.79; figure 5A), reaction times (main effect of block *β* = −0.003, CI = [−0.006, 0.011], *t*(116) = −0.66, *p* = 0.51; interaction *β* = −0.002, CI = [−0.014, 0.010], *t*(116) = −0.38, *p* = 0.70; figure 5B) or peak velocity (main effect of block *β* = −0.015, CI = [−0.066, 0.036], *t*(116) = −0.59, *p* = 0.56; interaction *β* = −0.024, CI = [−0.047, 0.096], *t*(116) = −0.67, *p* = 0.50; figure 5D). Finally, in line with the observed speed-accuracy functions, punishment context did affect radial accuracy, with accuracy increasing compared to the rewarding context (main effect of block, *β* = 0.10, CI = [0.019, 0.19], *t*(116) = 2.42, *p* = 0.017; figure 5F), although no interaction was observed (*β* = −0.07, CI = [−0.19, 0.05], *t*(116) = −1.16, *p* = 0.25). This can be directly observed when comparing baseline values, as radial error in the -0p condition was on average smaller than in the 0p condition (figure 5F, pink plot).

**Figure 5.**
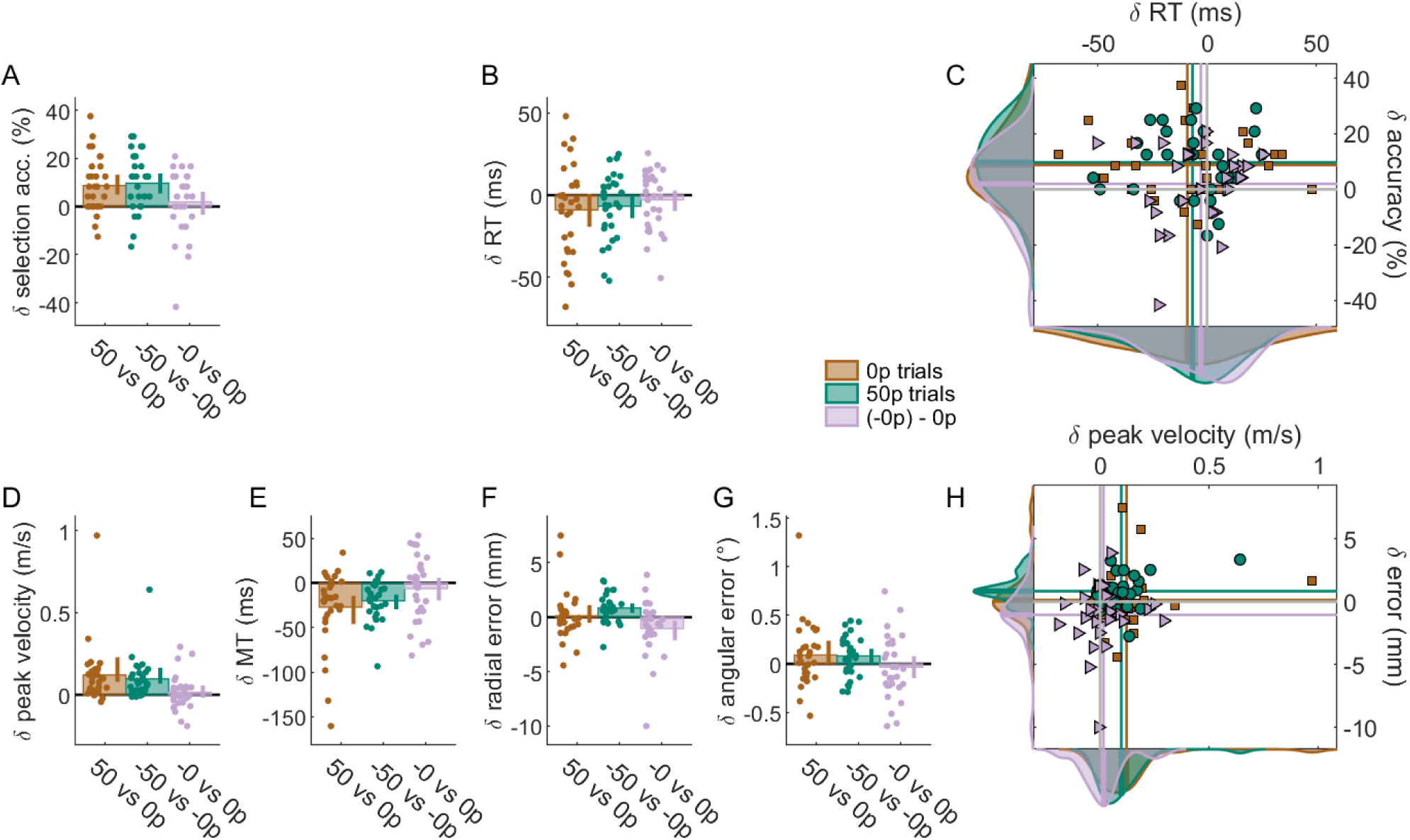
Reward and punishment have a similar effect on selection, but not on execution. For all bar plots, data was normalised to baseline performance (0p or -0p) for each individual. Bar height indicates group mean, dots represent individual values and error bars indicate bootstrapped 95% CIs of the mean. A. Selection accuracy. B. Mean reaction times for each participant. C. Scatterplot of mean reaction time against selection accuracy. Values are normalised to 0p trials. The coloured lines indicate mean values for each condition, and the solid grey lines indicate the origin, that is, 0p performance. Data distributions are displayed on the sides, with transversal bars indicating the mean of the distribution. Squares and triangles indicate +50p and (−0p)-0p trials, respectively. D. Mean peak velocity. E. Movement times. F. For radial error, punishment did not protect against an increase in error, while reward did. However, a difference can be observed between the baselines (blue bar). G. Angular error. H. Scatterplot showing execution speed (peak velocity) against execution accuracy (radial error), similar to C.

### 2.3 Reward reduces execution error through increased feedback correction and late noise resistance

How do reward and punishment lead to these improvements in motor performance? In saccades, it has been suggested that reward increases feedback control, allowing for more accurate end-point performance. To test for this possibility, we performed the same time-time correlation analysis as described in Manohar et al. (2019). Specifically, we assessed how much the set of positions at time *t* across all trials correlated with the set of positions at any other time *t± n, e.g. t* +1 or *t*− 5. If movements are stereotyped across trials, this correlation will be high because the early position will provide a large amount of information about the later or earlier position. On the other hand, if trajectories are variable over time within a trial, the correlation will decrease because there will be no consistency in the evolution of position over time. Importantly, the latter occurs with high online feedback because corrections are not stereotyped, but rather dependent on the random error on a given trial (Manohar et al., 2019). If the same mechanism is at play during reaching movements as in saccades, a similar decrease in time-time correlations should be observed.

All timepoints correlations were performed by comparing position over trials by centiles, leading to 100 timepoints along the trajectory (figure 6A-G). Across experiments 1 and 2, we observed an increase in time-time correlation in the late part of movement both with reward and punishment (figure 6H-K), although this did not reach significance in the 50p-0p condition of the second experiment (figure 6J) and was only marginally significant in the 10p-0p condition (figure 6H). In contrast, the early to middle part of movement showed a clear decorrelation that was significant in three conditions but not in the 50p-0p condition of the first experiment. Surprisingly, no difference was observed when comparing baseline trials from experiment 2 (figure 6L), which is at odds with the behavioural observations that radial error was reduced in the -0p condition compared to 0p (figure 5F). Overall, although quantitative differences are observed across cohorts, their underlying features are qualitatively similar (with the exception of the baselines contrast; figure 6L), displaying a decrease in correlation during movement followed by an increase in correlation at the end of movement. This suggests that a common mechanism may take place. To assess the global trend across cohorts, we pooled all cohorts together *a posteriori*, and indeed observed a weak early decorrelation, followed by a strong increase in correlation late in the movement (figure 6M). Interestingly, this consistent biphasic pattern across conditions and experiments is the opposite to the one observed in saccades (Manohar et al., 2019). Therefore, this analysis would suggest that reward/punishment causes a decrease in feedback control during the late part of reaching movements. However, a reduction in feedback control should result in a decrease in accuracy which was not observed in our data. A more likely possibility is that another mechanism is being implemented that enables movements to be performed with enhanced precision under reward and punishment.

**Figure 6.**
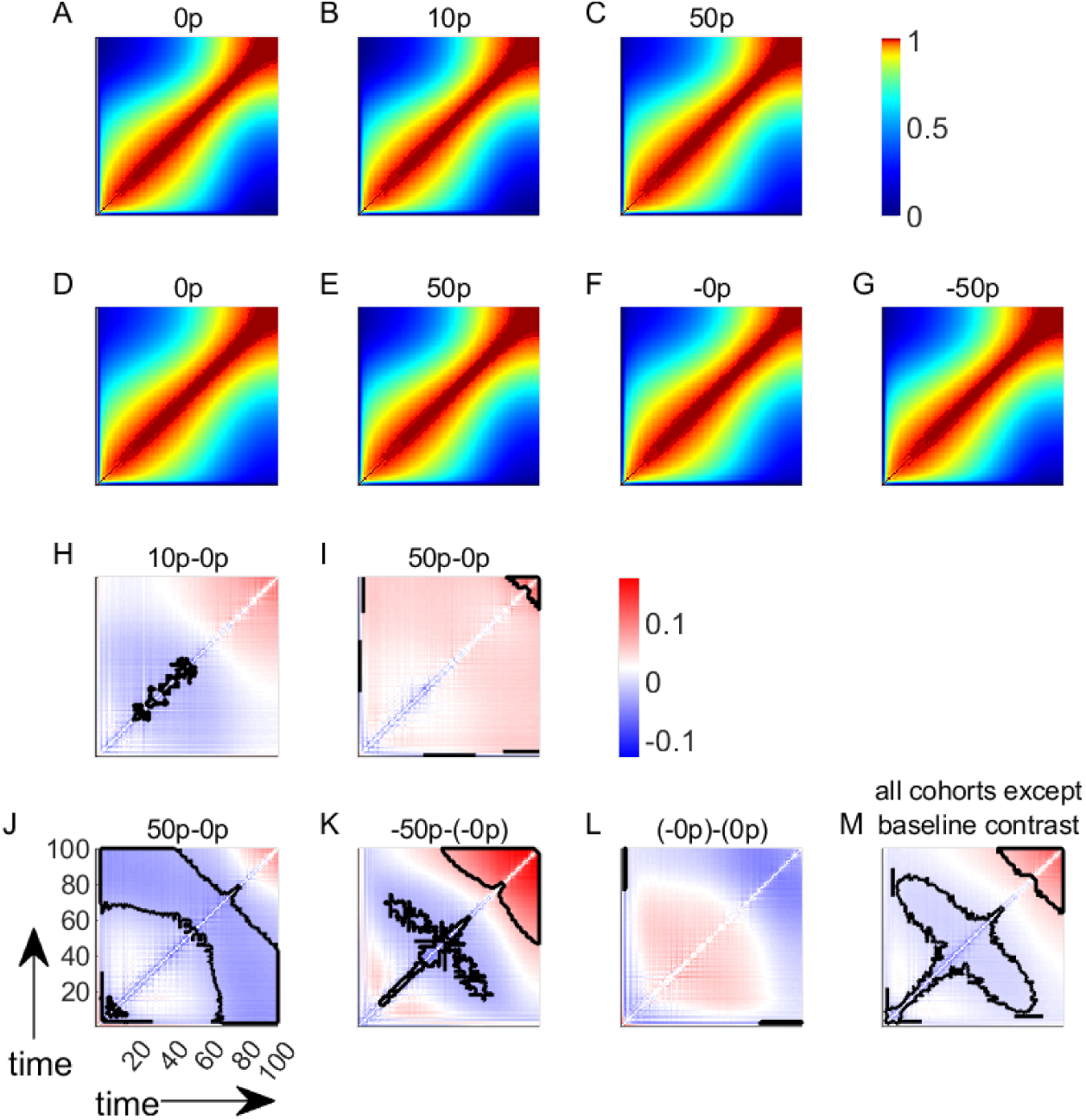
Time-time correlation maps show that monetary reward and punishment have a biphasic effect on the reach timecourse. A-C. Time-time correlation maps for all trial types (0p, 10p 50p) in Experiment 1. Colours represent Fisher-transformed Pearson correlation values. For each map, the lower left and upper right corners represent the start and the end of the reaching movement, respectively. Note that the colour maps are non-linear to enhance readability. D-G. Time-time correlation maps for all trial types (0p,50p,-0p,-50p) in Experiment 2. H-I. Comparison of fisher-transformed correlation maps with the respective baseline map (A) for Experiment 1. Clusters of significance after cluster-wise correction for multiple comparisons are indicated by a solid black line. J-L. Similar comparisons for Experiment 2, with each condition’s respective baseline (D and F). M. Similar comparison when pooling all contrasts except the baselines contrasts together.

One possible candidate is muscle co-contraction. By simultaneously contracting agonist and antagonist muscles around a given joint, the nervous system is able to regulate the stiffness of that joint. Although this is an extremely energy inefficient mechanism, it has been repeatedly shown that it is very effective at improving arm stability in the face of unstable environments such as force fields (Franklin et al., 2003). Critically, it is also capable of dampening noise (Selen et al., 2009), which arises with faster reaching movements, and therefore enables more accurate performance (Todorov, 2005). Therefore, it is possible that increased arm stiffness could, at least partially, underlie the effects of reward and punishment on motor performance.

### 2.4 Simulation of time-time correlation maps with a simplified dynamical system

To assess if the correlation maps we observed are in line with this interpretation, we performed simulations using a simplified control system (Manohar et al., 2019) and evaluated how it responded to hypothesised manipulations of the control system. Let us represent the reach as a discretised dynamical system (Todorov, 2004):

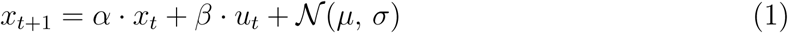

The state of the system at time *t* is represented as *x*_*t*_, the motor command as *u*_*t*_, and the system is susceptible to a random gaussian process with mean *µ* = 0 and variance *σ* = 1. *α* and *β* represent the environment dynamics and control parameter, respectively. For simplicity, we initially assume that *α* = 1, *β* = 0 and that *x*_0_ = 0. Therefore, any deviation from 0 is solely due to the noise term that contaminates the system at every time step.

We performed 1000 simulations, each including 1000 timesteps, and show the time-time correlation maps of the different controllers under consideration. First, we assume that no feedback has taken place (*β* = 0, equation 1). The system is therefore only driven by the noise term (figure 7A). The controller can reduce the amount of noise, *e.g.* through an increase in stiffness (Selen et al., 2009). This can be represented as *x*_*t*+1_ = *x*_*t*_ + *γ* · *𝒩* (*µ, σ*) with *γ* = 0.5. However, this would not alter the correlation map (figure 7B-C) as was previously shown (Manohar et al., 2019) because the noise reduction occurs uniformly over time. Now, if a feedback term is introduced with *β* = −0.002 and *u*_*t*_ = *x*_*t*_, the system includes a control term that will counter the noise and becomes:

**Figure 7.**
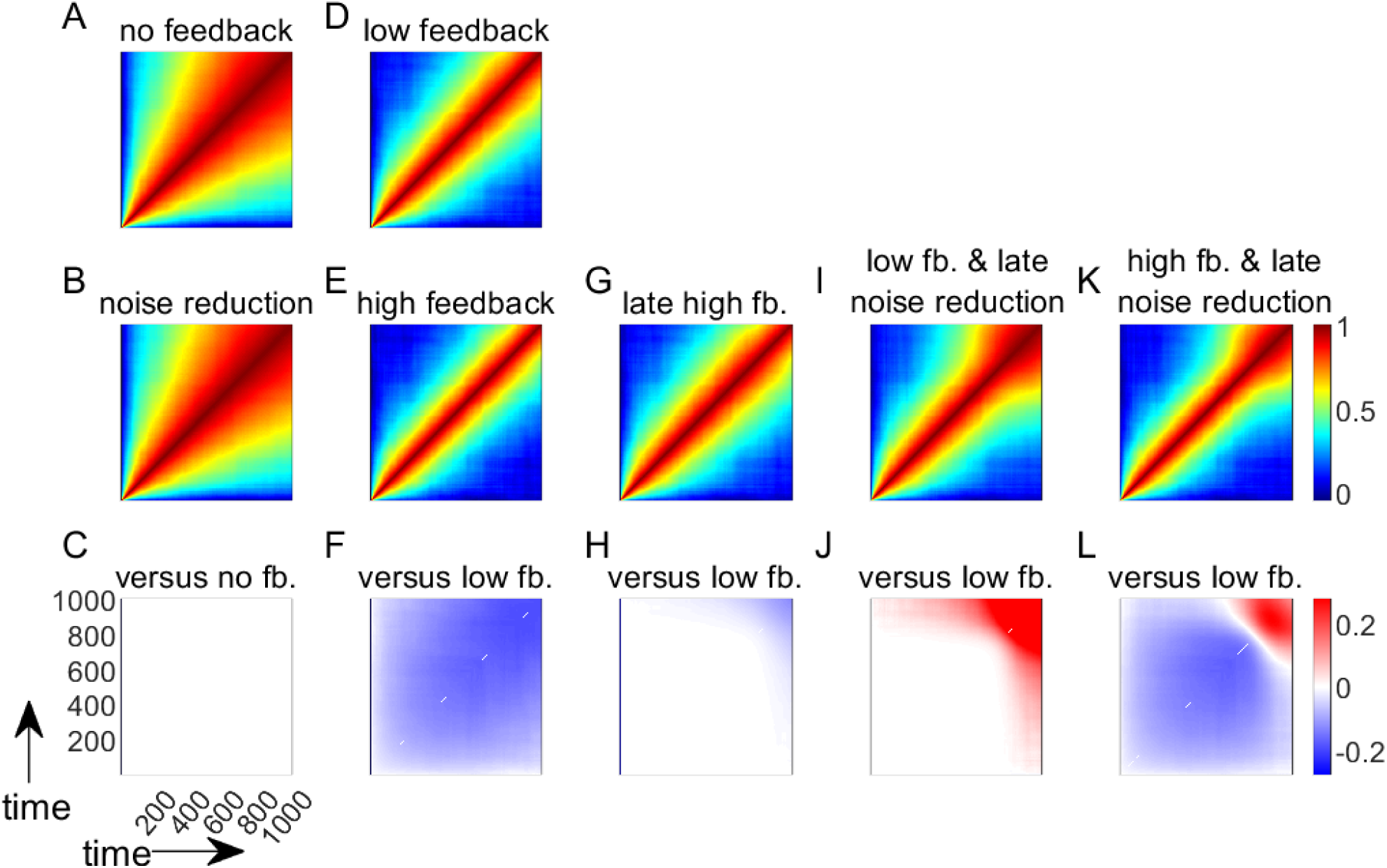
Simulations of time-time correlation map behaviour under different models of the reward- and punishment-based effects on motor execution. A,D. Time-time correlation maps of both control models. Colours represent Fisher-transformed Pearson correlation values. For each map, the lower left and upper right corners represent the start and the end of the reaching movement, respectively. B,E,G,I,K. Time-time correlation maps of plausible alternative models. C,F,H,J,L. Comparison of models with their respective baseline models.

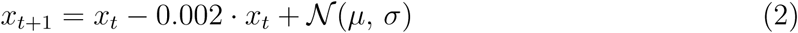

With such a corrective feedback term, the goal of the system becomes to maintain the state at 0 for the duration of the simulation. This is equivalent to assuming that *x* represents error over time and the controller has perfect knowledge of the optimal movement to be performed. Higher feedback control (*β* = −0.003) would reduce errors even further. Comparing this high feedback model with the low feedback model (equation 2; figure 7D-E), we see that the contrast (figure 7F) shows a reduction in time-time correlations similar to what is observed in the late part of saccades (Manohar et al., 2019) and in the early part of arm reaches in our dataset (figure 6H-K). Since our dataset displays a biphasic correlation map, it is likely that two phenomena occur at different timepoints during the reach. To simulate this, we altered the original model by including a sigmoidal step function *S*(*t*) that is inactive early on (*S*_0_ = 0) and becomes active 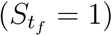 during the late part of the reach (see section Model simulations for details). This leads to two possible mechanisms, namely, a late increase in feedback or a late reduction in noise:

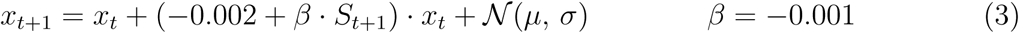

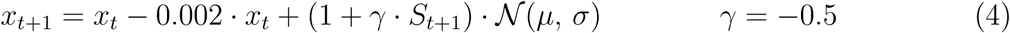

The results show that a late increase in feedback causes decorrelation at the end of movement (equation 3; figure 7G-H), which is the opposite of what we observe in our results. However, similar to our behavioural results, a late reduction in noise causes an increase in the correlation values at the end of movement (equation 4; figure 7I-J). Therefore, our results (figure 6H-K) appear to be qualitatively similar to a combined model in which reward and punishment cause a global increase in feedback control and a late reduction in noise (equation 5; figure 7K-L):

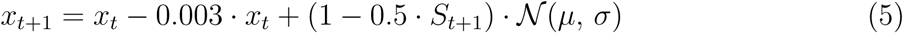

The simulations displayed here incorrectly assume that the noise term remains the same throughout the reach (Shadmehr & Krakauer, 2008; Todorov, 2004) and that feedback can account for errors from one timestep to the next, that is nearly immediately (Bhushan & Shadmehr, 1999). To explore if these features would alter our observations, we simulated two alternative sets of models. A first set included a bell-shaped noise term similar to a reach with signal-dependent noise under minimum jerk conditions (figure 7 supplement 1), and a second set included a delay of 400 timesteps in the feedback response (figure 7 supplement 2). Both sets of simulation produced results similar to those observed in the original set of models.

### 2.5 Quantitative model comparison

To formally test which candidate model best describes our empirical observations, we fitted each of them to the experimental datasets. Each of the five empirical conditions displayed in figure 6H-L was kept separate, each condition representing a cohort, and their fit assessed separately. While individually fitted models present several advantages over group-level analysis, it has been argued that the most reliable approach to determine the best-fit model is to assess its performance both on individual and group data and compare the outcomes (Cohen et al., 2008; Lewandowsky & Farrell, 2011) and we will therefore follow this approach. We included six candidate models in our analysis: noise reduction (one free parameter *γ*; figure 7C), increased feedback (one parameter *β*; figure 7E), late feedback (one parameter *β*; figure 7H), late noise reduction (one parameter *γ*; figure 7J), increased feedback with late noise reduction (two parameters *β* and *γ*; figure 7L) and an additional model with noise reduction and a late increase in feedback control (two parameters *β* and *γ*).

Individual-level analysis resulted in the increased feedback with late noise reduction model being selected by a strong majority of participants for each cohort (cohort 1-5: *χ*^2^ = [97.6, 76.8, 74.4, 116.8, 83.2], all *p <* 0.001, figure 8A), confirming qualitative predictions. The best-fit model for each participant was defined as the model bearing the lowest Bayesian information critetion (BIC; figure 8B). This allowed us to account for each model’s complexity, because the BIC penalises models with more free parameters. Of note, the “baselines” cohort displayed the highest BIC for all models considered. However, this should not be surprising, considering that this cohort is the only one that showed no significant trend in its contrast map (figure 6L). To confirm that the selected model is indeed the most parsimonious choice, we compared the individual-level outcome to a group-level outcome. Each candidate model was fit to all individual correlation maps at once, thereby allowing for each free parameter to take a single value per cohort. This is equivalent to assuming that the parameters are not random but rather fixed effects, allowing us to observe the population-level trend with higher certainty, though at the cost of ignoring its variability (Cohen et al., 2008; Lewandowsky & Farrell, 2011). Again, for every cohort except the baseline cohort, the model with lowest residuals sum of squares (figure 9A) and lowest BIC (figure 9B) was the increased feedback with late noise reduction model – though the increased feedback model BIC was marginally lower for the large-reward cohort (ΔBIC= 4) and therefore was a similarly good fit. Finally, fitting all non-baseline cohorts yielded the same result.

**Figure 8.**
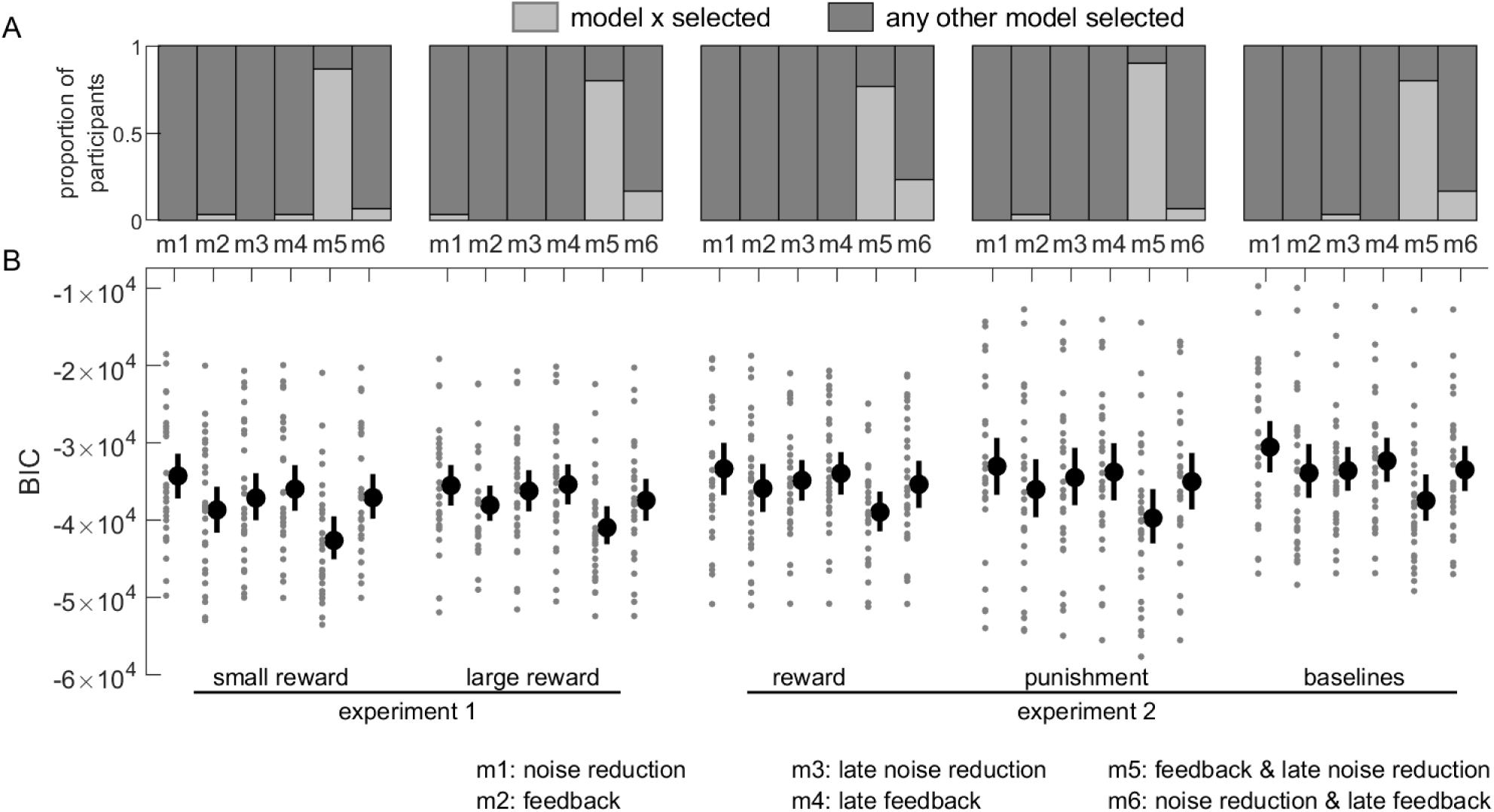
Model comparisons for individual fits. A. Proportion of participants whose winning model was the one considered (light gray) against all other models (dark gray) for every cohort. B. Individual and mean BIC values for each participant and each model. Lower BIC values indicate a better fit. Dots indicate individual BICs, the black dot indicates the group mean and the error bars indicate the bootstrapped 95% CIs of the mean. BIC: Bayesian information criterion.

**Figure 9.**
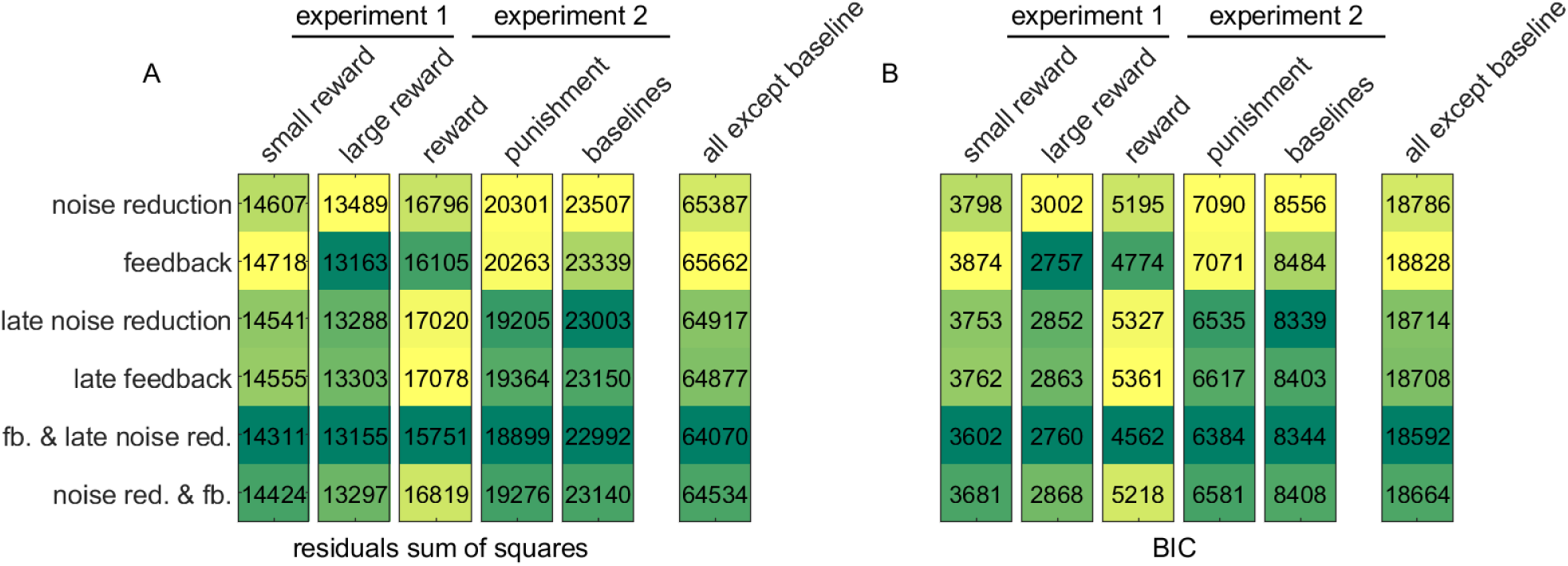
Model comparisons for group-level fits. A. residuals sum of squares for each model and cohort. Darker colours indicate lower values. B. Same as A for BIC. fb: feedback; noise red.: noise reduction; BIC: Bayesian information criterion.

Comparing group-level and invidividual-level model comparisons, we observe that the same model is consistently selected across all experimental cohorts besides the baselines cohort, corroborating the hypothesis that late noise reduction occurs alongside a global increase in feedback control in the presence of reward or punishment. One way to increase noise resistance during a motor task is by increasing joint stiffness, a possibility that we test in the following experiment.

### 2.6 The effect of reward on end-point stiffness at the end of the reaching movement

Next, we experimentally tested whether the reduction in noise observed in the late part of reward trials is associated with an increase in stiffness. For simplicity, we focused on the reward context only from this point. We recruited another set of participants (N=30) to reach towards a single target 20cm away from a central starting position in 0p and 50p conditions, and employed a well-established experimental approach to measure stiffness (Burdet et al., 2000; Selen et al., 2009). Specifically, during occasional “catch” trials (31% trials pseudorandomly interspersed) a fixed-length (8mm) displacement was applied to the robotic manipulandum immediately as participants stopped within the target. Because displacements of this amplitude were noticeable, participants were instructed to ignore them and not react, and we employed a low proportion of catch trials to reduce anticipation. The displacements were in 8 possible directions arrayed radially around the target (figure 10 supplement 1A). This displacement was transient, with a ramp-up, a plateau, and a ramp-down phase back to the original end-position. As the position was clamped during the plateau phase, velocity and acceleration were on average null, removing any influence of viscosity and inertia. Therefore, the amount of force required to maintain the displacement during plateau was linearly proportional to end-point stiffness of the arm (Perreault et al., 2002). The displacement profile of a participant is presented in figure 10 supplement 1B. Using a linear regression approach to fit the average recorded force during the plateau (grey area in figure 10 supplement 1B) against the displacement direction, we obtained the end-point stiffness matrices for all participants and all reward values. Stiffness matrices could then be visualised by plotting ellipses using the following equation:

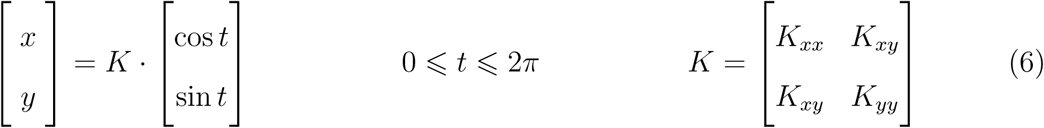

Because arm stiffness is strongly dependent on arm configuration, stiffness ellipses are usually oriented, with a long axis indicating a direction of higher stiffness (figure 10). This orientation is influenced by several factors, including position in Cartesian space (Mussa-Ivaldi et al., 1985). If reward affects stiffness as we hypothesised, the possibility that this effect is dependent on a target location must therefore be considered. To account for this, two groups of participants (N=15 per group) reached for a target 45° to the right or the left of the starting position.

**Figure 10.**
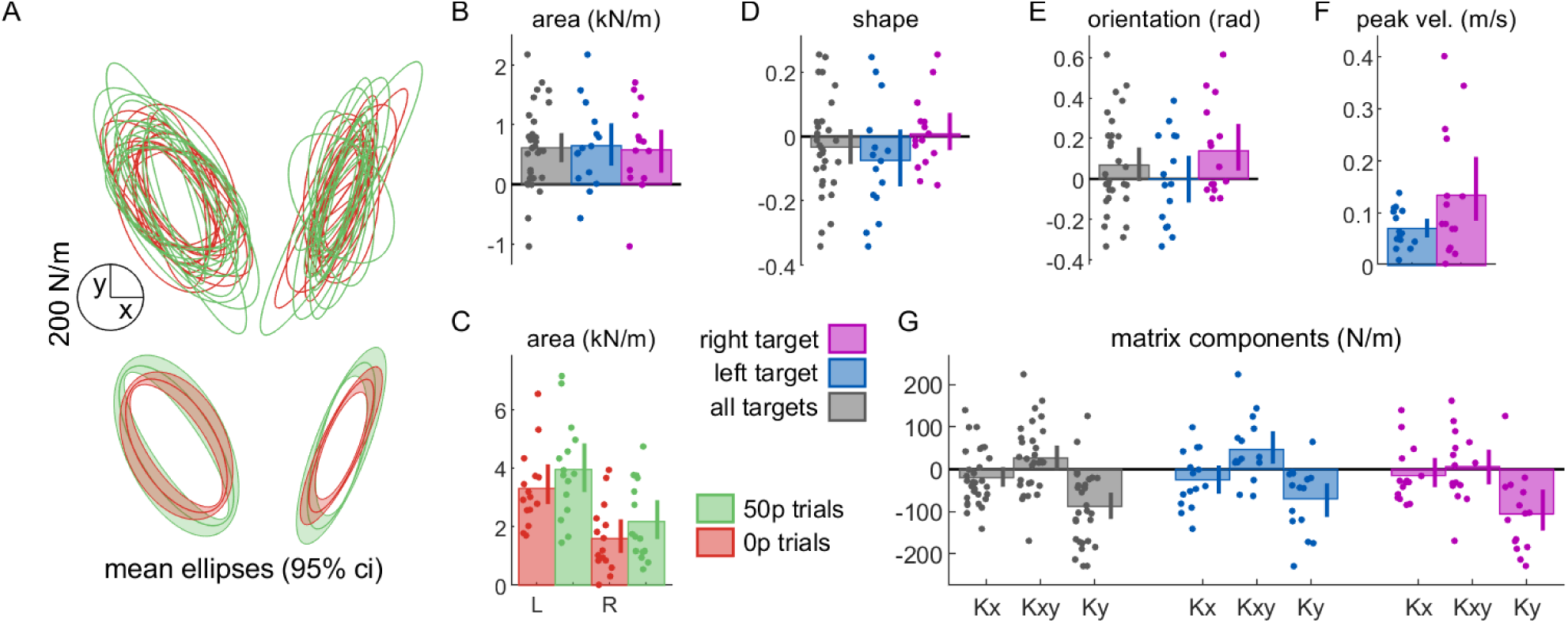
Reward increases stiffness at the end of movement. A. Individual (top) and mean (down) stiffness ellipses. Shaded areas around the ellipses represent bootstrapped 95% CIs. Right and left ellipses represent individual ellipses for the right and left target, respectively. B. Ellipses area normalised to 0p trials. Error bars represent bootstrapped 95% CIs. C. Non-normalised area values are also provided to illustrate the difference in absolute area as a function of target (L: left target, R: right target). D. Ellipse shapes normalised to 0p trials. Shapes are defined as the ratio of short to long diameter of the ellipse. E. Ellipse orientation normalised to 0p trials. Orientation is defined as the angle of the ellipse’s long diameter. F. Peak velocity normalised to 0p trials. Peak velocity increased with reward. G. Stiffness matrix elements for 50p trials normalised to the stiffness matrix for 0p trials.

To quantify the global amount of stiffness, we compared the ellipse area across conditions (figure 10A-C). In line with our hypothesis, the area substantially increased in rewarded trials compared to non-rewarded trials (figure 10A,B). This effect of reward was very consistent across both target positions (figure 10B), even though absolute stiffness was globally higher for the left target (figure 10C). On the other hand, other ellipse characteristics, such as shape and orientation (figure 10D,E) showed less sensitivity to reward. However, since reward also increased average velocity (figure 10F), in line with our previous results, perhaps this increase in stiffness is a response to higher velocity rather than reward. To avoid this confound, we fitted a mixed-effect linear model, allowing for individual intercepts and target position intercept, where variance in area could be explained both by reward and velocity: *area* ∼ 1 + *reward* + *peak velocity* + (1|*participant*) + (1|*target*). As expected, reward – but not peak velocity – could explain the variance in ellipse area (peak velocity: *p* = 0.31; reward: *p <* 0.001; table in figure 10 supplement 2), confirming that the presence of reward results in higher global stiffness at the end of the movement. In contrast, fitting a model with the same explanatory variables to the *Ky* component of the stiffness matrices, which showed the greatest sensitivity to reward compared to the other components (figure 10G) revealed that not only reward (*p <* 0.001, Bonferroni corrected) but also peak velocity (p=0.016, Bonferroni-corrected; table in figure 10 supplement 3) explained the observed variance (model: *Ky* ∼ 1 + *reward* + *peak velocity* + (1|*participant*) + (1|*target*)). In comparison, no significant effects were found to relate to the *Kx* component (reward: *p* = 0.14, peak velocity: *p* = 1, Bonferroni-corrected; *Kx* ∼ 1 + *reward* + *peak velocity* +(1|*participant*)+(1|*target*)).

Because interactions with nested elements cannot be compared directly using a mixed-effect linear model (Schielzeth & Nakagawa, 2013; Zuur et al., 2010; Harrison et al., 2018), we employed a repeated-measure ANOVA to compare the interaction between reward and target on stiffness. No interaction between reward and target location were observed on area (*F* (1) = 0.069, *p* = 0.79, partial *η*^2^ *<* 0.001; figure 10A,C).

We conclude that end-point stiffness is sensitive to both reward and velocity. However, the velocity-driven increase in stiffness is specific to the dimension that this velocity is directed toward, while the reward-driven increase in stiffness is non-directional, at least in our task. This is likely because our task does not distinguish direction of error (*i.e.* error in the *y* dimension is not more punishing than in the *x* dimension) and so error must be reduced in all dimensions (Selen et al., 2009).

### 2.7 Reward does not alter end-point stiffness at the start of the movement

Finally, the time-time correlation maps also suggest that the increase in stiffness should only occur at the end of the reaching movement, since the early and middle parts show an opposite effect (decorrelation). Therefore, an increase in end-point stiffness should not be present immediately before the reach. To test this, participants (N=20) reached to 2 targets positioned 20cm away and 45° to the left and right of the starting position. On occasional catch trials (31% trials), a displacement akin to the previous experiment occurred in one of 8 possible directions at the time normally corresponding to target onset but after the reward information had been displayed (figure 11 supplement 1). Because participants voluntarily moved into the starting position after it appeared, they had sufficient time to process the reward information. Unlike the previous experiment, reward and velocity in the subsequent reach had no impact on stiffness, either by area (reward: *p* = 0.35; peak velocity: *p* = 0.75, table in figure 11 supplement 2) or by the matrix component *K*_*y*_ (reward: *p* = 0.19; peak velocity: *p* = 0.45, table in figure 11 supplement 3), corroborating our interpretation of the correlation map (figure 11).

**Figure 11.**
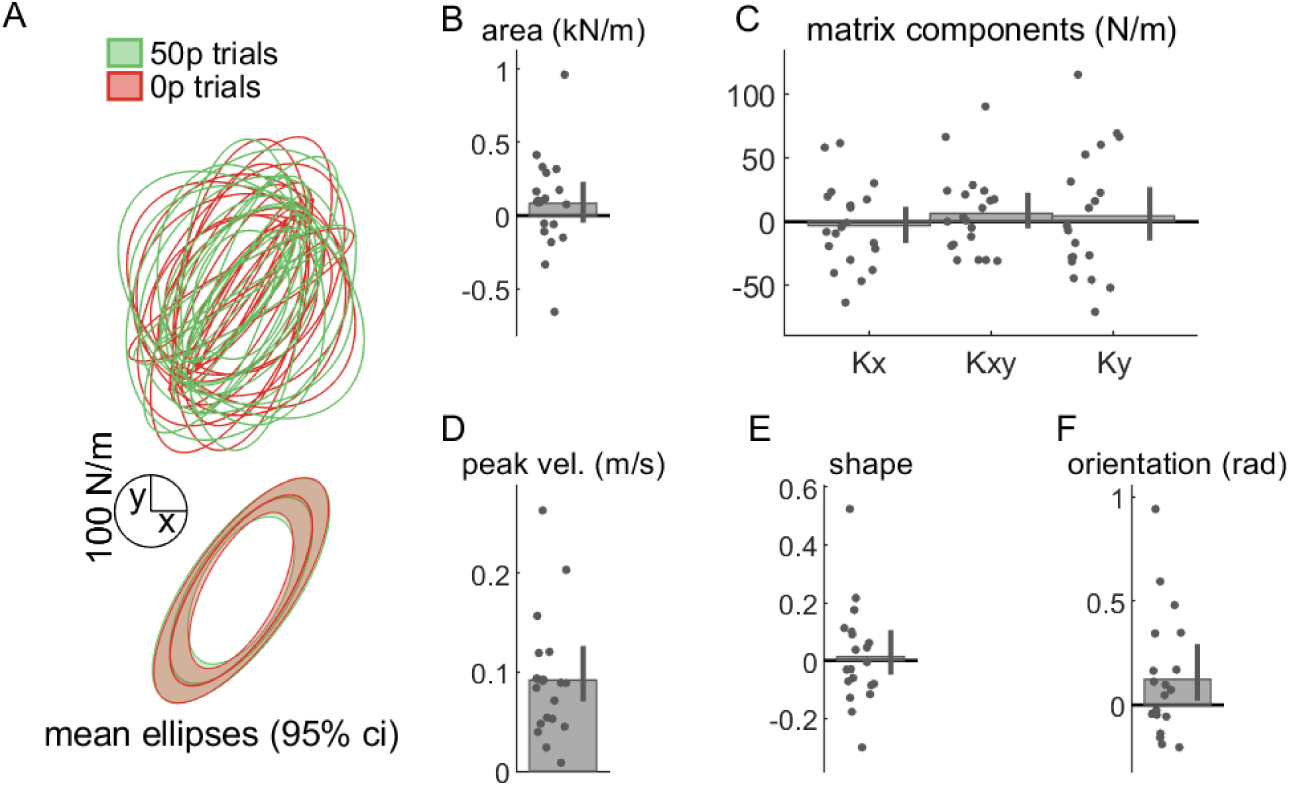
Reward does not alter stiffness at the start of movement. Individual (top) and mean (down) stiffness ellipses. Shaded areas around the ellipses represent bootstrapped 95% CIs. Right and left ellipses represent individual ellipses for the right and left target, respectively. Ellipses area normalised to 0p trials. Error bars represent bootstrapped 95% CIs. C. Stiffness matrix elements for 50p trials normalised to the stiffness matrix for 0p trials. D. Peak velocity normalised to 0p trials. E. Ellipse shapes normalised to 0p trials. Shapes are defined as the ratio of short to long diameter of the ellipse. F. Ellipse orientation normalised to 0p trials. Orientation is defined as the angle of the ellipse’s long diameter.

## 3 Discussion

In this study, we demonstrated that reward has the ability to simultaneously improve the selection and execution components of a reaching movement. Specifically, reward promoted the selection of the correct action in the presence of distractors, whilst also improving execution through increased speed and maintenance of accuracy. These results led to a shift in the speed-accuracy functions for both selection and execution. In addition, punishment had a similar impact on action selection and execution, although its impact was non-contingent for execution, in that it enhanced performance across all trials within a block, irrespective of the value of the current trial. Computational analysis revealed that the effect of reward on execution involved a combination of increased feedback control and noise reduction, which we then showed was due to an increase in arm stiffness at the end of the reaching movement – but not at the start of the movement. Overall, we confirm previous observations that feedback control increases with reward and offer a new error-managing mechanism that the control system employs under reward: regulation of arm stiffness.

Our results add to the previous literature arguing that reward increases execution speed in reaching (Chen, Holland & Galea, 2018; Pasquereau et al., 2007; Summerside et al., 2018) and saccades (Manohar et al., 2019, 2015; Takikawa et al., 2002). However, our results deviate from several reports in some respects. First, in a serial reaction time study, it was demonstrated that reward and punishment both reduced reaction times in humans (Wachter et al., 2009), while reaction times are not significantly altered by reward and punishment in our study. However, serial reaction time tasks strongly emphasise reaction times as a measure of learning independently of other variables, and interestingly, the authors show that punishment also led to a non-contingent effect on performance, while reward did not, similar to our results. A possible interpretation is that the motor system presents a similar bias to punishment to what is regularly reported in prospect theory and decision-making literature (**??**) – a phenomenon dubbed “loss aversion”. Next, radial accuracy has been shown to improve with reward, both in monkeys (Kojima & Soetedjo, 2017; Takikawa et al., 2002) and humans (Manohar et al., 2019, 2015), but these studies all focused on saccadic eye movements. In contrast, one reported case in a reaching task showed improvements in angular accuracy (Summerside et al., 2018). However, accuracy requirements in their noreward condition were minimal, possibly allowing for larger improvements to be expressed compared to our task, and potentially explaining why we did not observe similar improvement in radial or angular accuracy. Finally, while other studies have shown that speed-accuracy functions can shift with practice (Reis et al., 2009; Telgen et al., 2014), it is noteworthy that reward has a capacity to do so in what seems a nearly instantaneous time-scale, that is, from one trial to the next. Indeed, trials bearing different reward values were randomly intertwined in our study, meaning that this shift occurs within one trial. In contrast, the shift in speed-accuracy function observed with motor learning can take hours or even days to occur (Telgen et al., 2014).

### 3.1 Implications of increased stiffness with reward

While it is well established that stiffness has a beneficial effect on motor performance, our work provides the first set of evidence that this mechanism is employed in a rewarding context. Stiffness itself could be regulated through a change in co-contraction of antagonist muscles, which is a simple but costly method to increase stiffness and enhance performance against noise (Gribble et al., 2003; Selen et al., 2009; Ueyama & Miyashita, 2013; Ueyama et al., 2011). The presence of reward may make such cost “worthy” of the associated metabolic expense (Todorov, 2004), as has been shown in reaching in non-human primates (Ueyama & Miyashita, 2014). Another possibility is that the stretch reflex is increased, leading to a stronger counter-acting force produced against the perturbation. For instance, the stretch reflex is sensitive to cognitive factors such as postural threat (standing next to a signficant height; Horslen et al., 2018). Nevertheless, the contribution of stiffness in reward-based performance has implications for current lines of research on clinical rehabilitation that focus on improving rehabilitation procedures using reward (Goodman et al., 2014; Quattrocchi et al., 2017). While several studies report promising improvements, excessive stiffness may expose vulnerable clinical populations to increased risk of fatigue and even injury. Careful monitoring is therefore required to avoid this possibility.

### 3.2 Saccades and reaching movements differ in their utilization of stiffness control with reward

Contrary to our findings, previous work on saccades shows that reward had no effect on stiffness (Manohar et al., 2019). Therefore, our results demonstrate that reaching movements differ from saccadic control, in that it employs an additional error-managing mechanism. Why do saccadic and limb control employ dissociable control approaches?

A first explanation may be the difference in motor command profile. Saccadic control displays a remarkably stereotyped temporal pattern of activity, in which the saccade is initiated by a transient burst of action potentials from the motoneurons innervating the extraocular muscles (Joshua & Lisberger, 2015; Robinson, 1964). Critically, this burst of activity always reaches an output frequency close to its maximum nearly instantaneously in an all-or-nothing fashion (Joshua & Lisberger, 2015; Robinson, 1964), with only marginal variation based on reward and saccade amplitude (Manohar et al., 2019; Reppert et al., 2015; Robinson, 1964; Xu-Wilson et al., 2009). In comparison, motor commands triggering reaching movements present a great diversity of temporal profiles depending on task requirements, and often do not reach maximum stimulation level. This difference between the two controllers may result in a difference in the temporal pattern of motor unit recruitment. According to the size principle (Llewellyn et al., 2010), low-force producing, high-sensitivity motor units are always recruited first during a movement. However, those motor units are also more noisy due to their higher sensitivity (Dideriksen et al., 2012). Since saccades always rely on an all-or-nothing input pattern, all motor units may be quickly recruited, including high-force, low-sensitivity motor neurons that are normally recruited last. This would drastically reduce the production of peripheral noise, thus making co-contraction unnecessary (Dideriksen et al., 2012). This is in line with previous work showing peripheral noise has a minimal contribution to overall error in eye movements (Van Gisbergen et al., 1981) compared to internally generated noise (Manohar et al., 2019). Interestingly, evidence of the opposite has been reported for reaching, suggesting that execution rather than planning noise is dominant in reaching errors (van Beers et al., 2004). These dissociable activation patterns of motor commands could potentially explain the differences in error-managing mechanisms between saccadic control and reaching.

A second possibility is that the muscles considered in saccade and reaching have different size and innervation density. Although eyes muscles are smaller, they are remarkably more innervated than most peripheral skeletal muscles (Floeter, 2010; Porter et al., 1995) such as arm muscles recruited for reaching, leading to a greater quantity of motor units. Interestingly, it has been shown that motor noise arising at the muscle level scales negatively with the number of motor units in that muscle (Hamilton et al., 2004). This may lead to reduced levels of execution noise for eye movements compared to reaching movements, making stiffness regulation less necessary for saccades. However, this falsely assumes that the physiology of motor units in extraocular muscles is the same as in limb muscles (Buchthal & Schmalbruch, 1980), and so this last interpretation should be considered with care.

### 3.3 Increased feedback control and reward

It is less clear what kind of feedback control may play a role in reward-driven improvements. Feedback control encompasses several processes that share the aim of tracking of deviation from a motor plan to correct for it, with varying amount of delay to allow for travelling from the peripheral sensory receptors to the brain. This includes the spinal stretch reflex (∼25ms delay; Weiler et al., 2019), transcortical feedback (∼50ms; Pruszynski et al., 2011) and visual feedback (∼170ms for fast involuntary visual feedback responses; Carroll et al., 2019). While spinal stretch reflex is extremely fast, it is difficult to assume an effect of reward or motivation occurring at the spinal level. On the other hand, transcortical feedback includes primary motor cortex processing (Pruszynski et al., 2011), a structure that shows sensitivity to reward (Bundt et al., 2016; Galaro et al., 2019; Thabit et al., 2011). Consequently, an exciting possibility for future research is that transcortical feedback gain is directly enhanced by the presence of reward. Indirect evidence suggests that this may be the case, as feedback control of matching timescales is sensitive to urgency in reaching (Crevecoeur et al., 2013). This suggests that transcortical feedback gains can also be pre-computed before movement initiation to meet task demands. Finally, recent work shows that reward can indeed modulate visual feedback control in reaching (Carroll et al., 2019) at timescales of 170-220ms after movement onset, much faster than usually considered for this type of feedback control (Carroll et al., 2019; Kasuga et al., 2015). Despite this remarkable speed, considering our typical movement times, this would imply that feedback control is increased only after about half of the movement. Therefore, a more conservative possibility is that both transcortical and visual feedback gains increase in the presence of reward, though the former remains to be proved empirically.

In saccades, it has been shown that the feedback controller that underlies reward-driven improvements is located further upstream, at the movement computation stage. Indeed, although saccadic control is ballistic and therefore feedforward, the cerebellum can provide some form of feedback to adjust the end part of a saccade trajectory based on errors in the forward model prediction (Robinson, 1981; Chen-Harris et al., 2008; Frens & Donchin, 2009). More recently, Manohar et al. (2019) demonstrate that it is this feedback loop that accounts for the observed improvements in feedback control during saccades. Interestingly, evidence in humans show that cerebellar forward models do contribute to feedback control in reaching to compensate for sensory delays (Miall et al., 2007), and more recently, optogenetics manipulation in mice confirmed its involvement in enhancing end-point precision based on reaching kinematics (Becker & Person, 2019). Therefore, it is possible that reward also enhances this feedback loop, though this would only contribute to reducing noise arising at the higher, computational stage rather than at effector stage (Manohar et al., 2019). Furthermore, it should be noted that both in saccadic and reaching tasks, empirical evidence shows this form of feedback contributes exclusively during the last portion of the movement, which is in contradiction with what we observe here.

### 3.4 Limitations of the model

The model we employ presents several assumptions and limitations. First, it reduces the movement to errors over time, because it only deals with the deviation from zero. This is similar to assuming that a perfect knowledge of the movement to be performed is already acquired, because deviations are only a function of the noise term. Furthermore, since the model is concerned with maintaining the system at a given value rather than “travelling” to a novel position, the expected bell-shaped profile of motor commands (Shadmehr & Krakauer, 2008; Todorov, 2004) is abstracted away, and thus the noise term is not signal-dependent (Todorov, 2005). However, additional simulations show that adding a bell-shaped noise term does not qualitatively alter the observations of the original set of models. Furthermore, these simplifications can be overlooked when considering model selection, because it is only concerned about a directional change from an arbitrary control model (*i.e.* increase versus decrease in time-time correlation). However, it may impede reliable parameter estimation because it remains an abstraction that excludes particular features such as two-dimensional reaches or signal-dependent noise. Finally, noise can arise from different sources (*e.g.* planning noise, execution noise and sensory noise) with a different impact on the final reaching behaviour measured (Dhawale et al., 2017). Future work using simulations based on a more complete model of the arm may provide further information regarding the evolution of saccadic and reaching profiles over time and allow reliable parameter estimation.

### 3.5 Conclusion

In this study, we show that reward can improve the selection and execution components of reaching movement simultaneously. While we confirm previous suggestions that enhanced feedback control contributes to this improvement, we introduce a novel, peripheral rather than central mechanism by showing that global end-point stiffness is regulated by the monetary value of a given trial. Therefore, reward drives multiple error-reduction mechanisms which enable individuals to invigorate motor performance without compromising accuracy.

## 4 Methods

### 4.1 Participants

30 participants (2 males, median age: 19, range: 18-31) took part in experiment 1. 30 participants (4 males, median age: 20.5, range: 18-30) took part in experiment 2. 30 participants (10 male, median age: 19.5, range: 18-32) took part in experiment 3, randomly divided into two groups of 15. 20 participants (2 male, median age: 19, range: 18-20) took part in experiment 4. All participants were recruited on a voluntary basis and were rewarded with money (£7.5/h) or research credits depending on their choice. Participants were all free of visual (including colour discrimination), psychological or motor impairments. All the experiments were conducted in accordance with the local research ethics committee of the University of Birmingham, UK.

### 4.2 Task design

Participants performed the task on an end-point KINARM (BKIN Technologies, Ontario, Canada). They held a robotic handle that could move freely on a plane surface in front of them, with the handle and their hand hidden by a panel (figure 1A). The panel included a mirror that reflected a screen above it, and participants performed the task by looking at the reflection of the screen, which appeared at the level of the hidden hand. The sampling rate was 1kHz.

Each trial started with the robot handle bringing participants 4cm in front of a fixed starting position, except for experiments 3-4 to avoid interference with the perturbations during catch trials. A 2cm diameter starting position (angular size 3.15°) then appeared, bearing a colour that matched one of several possible reward values, depending on the experiment. The reward value was also displayed in 2cm-heigh text (angular size *∼*3.19°) under the starting position (figure 1C-D). Because colour luminance can affect salience and therefore detectability, luminance-adjusted colours were employed (see http://www.hsluv.org/). The colours employed were, in red-green-blue format, [76,133,50] (green), [217,54,104] (pink) and [59,125,171] (blue) for 0, 10 and 50p, respectively, and distractor colours were either green, pink or blue. To ensure that a specific colour did not bias the amount of distracted trials, we fitted a mixed-effect model *distracted ∼ colour* +(1 |*participant*)+(1|*reward*) with *colour* a 3-level categorical variable encoding the colour of the distractor target. Distractor colour did not explain any variance in selection error (*p* = 1.72 × 10^−69^, *p* = 0.46 and *p* = 0.82 for the intercept, pink and blue colours, respectively) confirming that the observed effect was not driven by distractor colours. From 500 to 700ms after participants entered the starting position (on average 587 *±* 354ms after the starting position appeared), a 2cm target (angular size ∼2.48°) appeared 20cm away from the starting position, bearing the same colour as the starting position. Participants were instructed to move as fast as they could towards it and stop in it. They were informed that a combination of their reaction time and movement time defined how much money they would receive, and that this amount accumulated across the experiment. They were also informed that end-position was not factored in as long as they were within 4cm of the target centre.

The reward function was a close-loop design that incorporated the recent history of performance, to ensure that participants received similar amounts of reward, and that the task remained consistently challenging over the experiment (Manohar et al., 2015; Reppert et al., 2018). To that end, the reward function was defined as:

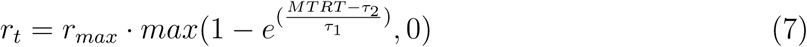

where *r*_*max*_ was the maximum reward value for a given trial, *MTRT* the sum of reaction time and movement time, and *τ*_1_ and *τ*_2_ adaptable parameters varying as a function of performance (figure 1B). Specifically, *τ*_1_ and *τ*_2_ were the median of the last 20 trials’ 3-4th and 16-17th fastest MTRTs, respectively, and were initialised as 400 and 800ms at the start of each participant training block. *τ* values were constrained so that *τ*_1_ *< τ*_2_ *<* 900 is always true. In practice, all reward values were rounded up (or down in the punishment condition of experiment 2) to the next penny so that only integer penny values would be displayed.

Targets were always of the same colour as the starting position (figure 1C). However, in experiments 1-2, occasional distractor targets appeared, bearing a different colour than the starting position (green, pink or blue depending on the correct target’s colour; figure 1D). Participants were informed to ignore these targets and wait for the second target to appear. Failure to comply in rewarded and punished trials resulted in no gains for this trial and an increase in loss by a factor of 1.2, respectively. The first target (distractor or not) appeared 500-700ms after entering the starting position using a uniform random distribution, and correct targets in distractor trials appeared 300-600ms after the distractor target using the same distribution.

When reaching movement velocity passed below a 0.3 m/s threshold, the end position was recorded, and monetary gains were indicated at the centre of the workspace. After 500ms, the robotic arm then brought the participant’s hand back to the initial position 4cm before the starting position.

In every experiment, participants were first exposed to a training block, where all targets had the same reward value equal to the mean of all value combinations used later in the experiment (*e.g.* if the experiment had 0p and 50p trials, the training reward amounted to 25p per trial). Participants were informed that money obtained during the training will not count toward the final amount they would receive. Starting position and target colours were all grey during training. The *τ* values obtained at the end of training were then used as initial values for the actual task.

### 4.3 Experimental design

#### 4.3.1 Experiment 1: reward-magnitude

There were 4 possible target locations positioned every 45° around the midline of the workspace, resulting in a 135° span (figure 1A). Participants first practiced the task in a 48-trial training block. They then experienced a short block (24 trials) with no distractors, and then a main block of 168 trials (72 distractors, 42.86%). Trials were randomly shuffled within each block. Reward values used during the task were 0, 10 and 50p.

#### 4.3.2 Experiment 2: reward-punishment

The same 4 target positions were used as experiment 1, and participants first practiced the task in a 48 trials training block. Participants then performed a no-distractor block and a distractor block (12 and 112 trials) in a rewarded condition (0p and 50p trials) and then in a punishment condition (−0p and -50p trials), in a counterbalanced fashion across participants. In the distractor blocks, 48 trials were distractor trials (42.86%). Before the punishment blocks, participants were told that they would start with £11 and that the slower they moved, the more money they lost. This resulted in participants gaining on average a similar amount of money on the reward and punishment blocks. They were also informed that if they missed the target or went to the distractor target, their losses on that trial would be multiplied by a factor of 1.2. The reward function was biased so that:

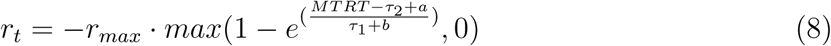

With *a* = 268.5 and *b* = −71.4. The update rule was also altered, with *τ*_1_ and *τ*_2_ the median of the last 20 trials’ 15-16th and 17-18th fastest MTRTs, respectively. These changes were obtained by fitting the performance data of the reward-magnitude experiment to a punishment function with free *a* and *b* parameters and free updating indexes to minimise the difference in average losses compared to the average gains observed in the reward-magnitude experiment. On average, participants gained £5.40 in the reward condition and lost £5.63 in the punishment condition (paired t-test: *t*(29) = −0.55, *p* = 0.58, *d* = −0.1; figure 5 supplement 2).

#### 4.3.3 Experiment 3: end-reach stiffness

In this task, each of two groups reached to a target located 20cm from the starting position, at +45 and −45° from the midline for the first and second group, respectively. On occasional catch trials, when movement velocity passed under a 0.3m/s threshold, a 300ms-long, 8mm displacement pushed participants away from their starting position and back, allowing us to measure end-point stiffness (see section Data analysis and figure 10 supplement 1). No distractor trials were employed in this experiment.

Participants performed two training sessions, one with no catch trials (25 trials) and one with 4 catch trials out of 8 trials, in four possible directions from 0 to 270° around the end position to familiarise participants with the displacement. Participants then performed the main block with 64 catch trials out of 200 trials (32%) and 0p and 50p reward values. During the main block, displacements were in 1 of 8 randomly assigned directions from 0-315° around the end-position (figure 10 supplement 1A). We used sessions of 233 trials to ensure session durations remained short, ruling out any effect of fatigue on stiffness as co-contraction is metabolically taxing. To ensure that any measure of stiffness was not due to differences in grip position or a loose finger grip, participant’s hands were restricted with a solid plastic piece which held the wrist straight and a reinforced glove that securely strapped the fingers around the handle during the entire task.

#### 4.3.4 Experiment 4: start-reach stiffness

The experiment was identical to experiment 3, except that the catch trials occurred in the start position (figure 11 supplement 1A) at the time the target was supposed to appear. To ensure participants remained in the starting position, two different targets (±45° from midline) were used to maintain directional uncertainty. Participants had 24 trials during the no-catch-trial training, 16 trials during the catch-trial training (8 catch trials), and 200 trials during the main block, with 64 (32%) catch trials. Displacements always occured 500ms after entering the starting position, to avoid a jitter-induced bias in stiffness measurement. In non-catch trials, targets also appeared after a fixed delay of 500ms.

### 4.4 Data analysis

All the analysis code is available on the *Open Science Framework* website, alongside the experimental datasets at https://osf.io/7as8g/. Analyses were all made in Matlab (Mathworks, Natick, MA) using custom-made scripts and functions.

Trials were manually classified as distracted or non-distracted. Trials that did not include a distractor target were all considered non-distracted. Distracted trials were defined as trials where a distractor target was displayed, and participants initiated their movement (*i.e.* exited the starting position) toward the distractor instead of the correct target. If participants readjusted their reach “mid-flight” to the correct target or initiated their movement to the right target and readjusted their reach to the distractor, this was still considered a distracted trial. On very rare occasions (*<*20 trials in the whole study), participants exited the starting position away from the distractor but before the correct target appeared; these trials were not considered distracted.

Reaction times were measured as the time between the correct target onset and when the participant’s distance from the centre of the starting position exceeded 2cm. In trials that were marked as “distracted” (*i.e.* participant initially went to the distractor target), the distractor target onset was used. In distractor-bearing trials, the second target did not require any selection process to be made, as the appearance of the distractor informed participants that the next target would be the right one. For this reason, reaction times were biased toward a faster range in trials in which a distractor target appeared, but participants were not distracted by it. Consequently, mean reaction times were obtained by including only trials with no distractor, and trials with a distractor in which participants were distracted. For the same reason, trials in the first block were not included because no distractor was present, and no selection was necessary. For every other summary variable, we included all trials that were not distracted trials, including those in the first block.

In experiments 1-2, we removed trials with reaction times higher than 1000ms or less than 200ms, and for non-distracted trials we also removed trials with radial errors higher than 6cm or angular errors higher than 20°. Overall, this resulted in 0.3% and 0.7% trials being removed from experiment 1 and 2, respectively. Speed-accuracy functions were obtained for each participant by binning data in the *x*-dimension into 50 quantiles and averaging all *y*-dimension values in a *x*-dimension sliding window of a 30-centile width (Manohar et al., 2015). Then, each individual speed-accuracy function was averaged by quantile across participants in both the *x* and *y* dimension.

Time-time correlation analyses were performed exclusively on non-distracted trials. Trajectories were taken from exiting the starting position to when velocity fell below 0.1m/s. They were rotated so that the target appeared directly in front of the starting position, and *y*-dimension positions were then linearly interpolated to a hundred evenly spaced timepoints. We focused on the *y* dimensions because it displays most of the variance (figure 12). Correlation values were obtained on *y*-positions and fisher-transformed before follow-up analyses (Manohar et al., 2019).

**Figure 12.**
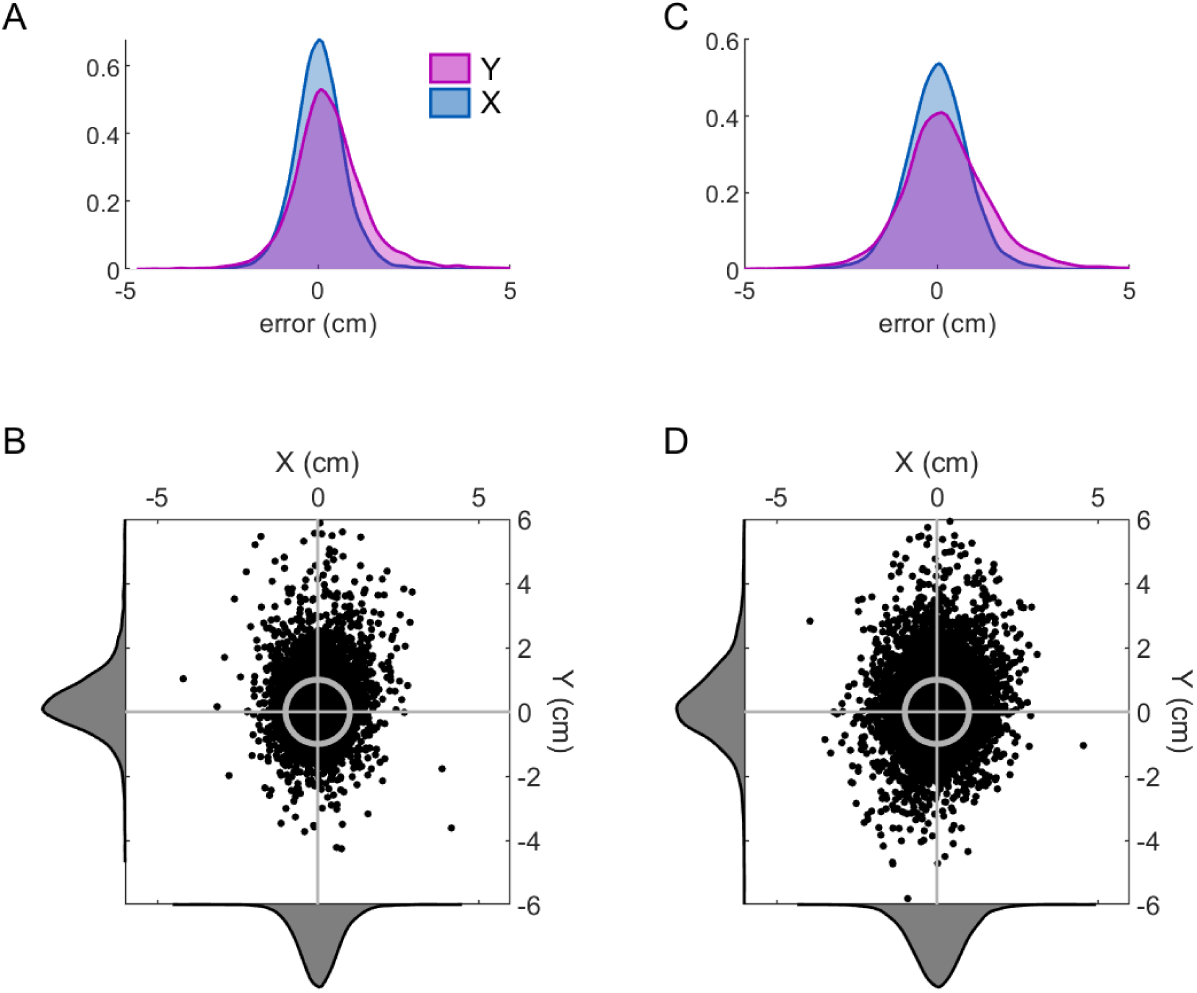
Distribution of errors at the end of the reach in the *x* and *y* dimension. A. Density function of errors in the *x* and *y* dimensions for experiment 1. B. Scatterplot of *x* versus *y* error after rotation of all target locations to a frontal location. The horizontal and vertical grey lines indicate the centre of the target, and the circle indicates its size. Density distributions can be observed on the sides. C-D. Same as A-B for experiment 2

For experiments 3-4, positions and servo forces in the *x* and *y* dimensions between 140-200ms after perturbation onset were averaged over time for each catch trial (Franklin et al., 2003; Selen et al., 2009). Then, the stiffness values were obtained using multiple linear regressions (function *fitlm* in Matlab). Specifically, for each participant, *K*_*xx*_ and 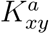 were the resulting *x* and *y* coefficients of *F*_*x*_ ∼ 1 + *x* + *y* and 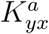 and *K*_*yy*_ were the resulting *x* and *y* coefficients of *F*_*y*_ ∼ 1 + *X* + *Y*. Data points whose residual was more than 3 times the standard error of all residuals were excluded (1.56% and 2.27% for experiment 3 and 4, respectively). Then, we can define the asymmetrical stiffness matrix:

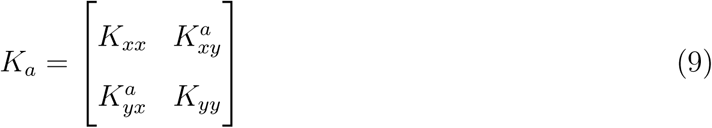

And the symmetrical stiffness matrix that we will use in subsequent analysis:

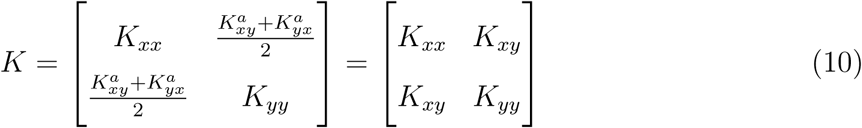

These matrices can be projected in Cartesian space using a sinusoidal transform (equation 6), resulting in an ellipse. This ellipse can be characterised by its shape, orientation and ratio, which we obtained using a previously described method (Perreault et al., 2002).

#### 4.5 Statistical analysis

Although for most experiments we employed mixed-effect linear models to allow for individual intercepts, we used a repeated-measure ANOVA in experiment 1 to compare each reward magnitudes against each other independently. This allowed us to assess the effect of reward without assuming a magnitude-scaled effect in the first place. Paired-sample t-tests were used when one-way repeated-measure ANOVA reported significant effects, and effect sizes were obtained using partial *η*^2^ and the Cohen’s d method. For experiment 2, we used mixed-effect linear models. For experiments 3 and 4, mixed-effect linear models were also used to account for a possible confound between reward and peak velocity in stiffness regulation, while accounting for individual differences in speed using individual intercepts. Since experiment 3 included a nested design (*i.e.* participants were assigned either to the right or left target but not both), we tested for an interaction using a two-way mixed-effect ANOVA to avoid an artificial inflation of p-values (Zuur, 2009). For all ANOVA, Bonferroni corrections were applied where appropriate, and post-hoc paired-sample t-tests were used if ANOVA produced significant results. Bootstrapped 95% confidence interval of the mean were also obtained and plotted for every group.

Since trials consisted of straight movements toward the target, we considered position in the *y* dimension – *i.e.* radial distance from the starting position – to obtain time-time correlation maps because it expresses most of the variability. To confirm this, reach trajectories were rotated so the target was always located directly in front, and error distribution in the *x* and *y* dimension was compared for both experiment 1 (figure 12A-B) and 2 (figure 12C-D). The *y* dimension indeed displayed a larger spread in error (experiment 1: *t*(11156) = −16.15, *p <* 0.001, *d* = −0.31; experiment 2: *t*(14852) = −13.68, *p <* 0.001, *d* = −0.22). Time-time correlation maps were analysed by fitting a mixed-linear model for each timepoint (Manohar et al., 2019; Zuur, 2009) allowing for individual intercepts using the model *z* ∼ *reward* + (1|*participant*), with *z* the fisher-transformed Pearson coefficient *ρ* for that timepoint. Then clusters of significance, defined as timepoints with p-values for reward of less than 0.05, were corrected for multiple comparisons using a cluster-wise correction and 10,000 permutations (Maris & Oostenveld, 2007; Nichols & Holmes, 2002). This approach avoids unnecessarily stringent corrections such as Bonferroni correction by taking advantage of the spatial organisation of the time-time correlation maps (Maris & Oostenveld, 2007; Nichols & Holmes, 2002).

### 4.6 Model simulations

The simulation code is available online on the *Open Science Framework* URL provided above. Simulation results were obtained by running 1000 simulations and obtaining time-time correlation values across those simulations. The sigmoidal step function *S*(*t*) used for simulations of the late component was a Gaussian cumulative distribution function such as:

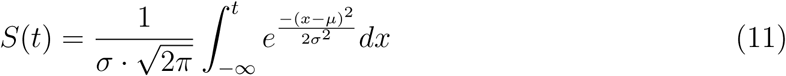

with *σ* = 0.5, *µ* = 0.8 (or 800 for a 1000 timesteps simulation) and *t*_0_ *< t < t*_*f*_ is the simulation timestep. It should be noted that the use of a sigmoidal function is arbitrary and may be replaced by any other step function, though this will only alter the simulation outcomes quantitatively rather than qualitatively. Values of the feedback control term are taken from Manohar et al. (2019). On the other hand, different noise terms were taken for our simulations because previous work only manipulated one parameter per comparison, whereas we manipulated both noise and feedback at the same time in several models (equations 4 and 5) and the model is more sensitive to feedback control manipulation than to noise term manipulation.

Two alternative sets of models were used to assess the effect of signal-dependent noise and delay in feedback corrections, respectively. For the first set, the noise term was redefined as *𝒩 (µ, σ* (*t*)) with:

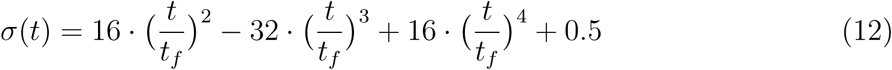

with equation 12 being proportional to the velocity profile of a minimum jerk reaching movement (Flash & Hogan, 1985). Here, the equation was adjusted so that 0.5 ⩽ *σ* (*t*) ⩽ 1.5, *σ* (0) = *σ*(*t*_*f*_) = 0.5 and *σ* (*t*_*f*_ */*2) = 1.5. The second set of models included a delay in feedback corrections, so that the feedback term *β* · *x*_*t*_ and its equivalent in different model variations became *β* · *x*_*t*−399_. A four hundred timesteps delay was chosen because observed movement times in the reward-magnitude and reward-punishment experiments were on average between 350-400ms (figure 3 supplement 1E and figure 5 supplement 1E), resulting in a feedback delay of ∼ 350 × 400*/*1000 = 140ms, which is within the range of feedback control delays expressed during reaching tasks (Pruszynski et al., 2011; Carroll et al., 2019).

Regarding model selection, comparisons were performed by fitting each of five datasets to six candidate models:

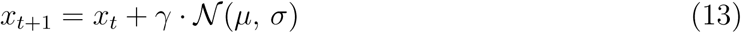

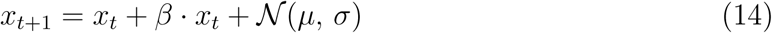

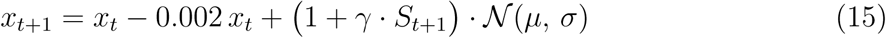

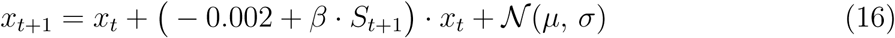

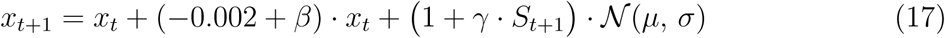

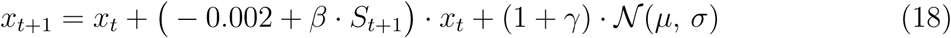

with equation 13 representing a model with noise reduction, equation 14 a model with increased feedback control, equation 15 a model with late noise reduction, equation 16 a model with late increase in feedback control, equation 17 a model with increased feedback and late noise reduction and equation 18 a model with late noise reduction and increased feedback. The free parameters were *β* and *γ*, with the last two model including both of them and all the others including one, according to the equations. *S*(*t*) was a step function as indicated in equation 11 and was fixed. 1000 simulations were done with 100 timesteps per simulation. Time-time correlation maps were then fisher-transformed and substracted to a control model *x*_*t*+1_ = *x*_*t*_ + *𝒩* (*µ, σ*) for equation 13 and *x*_*t*+1_ = *x*_*t*_ −0.002·*x*_*t*_ + *𝒩* (*µ, σ*) for all other models to obtain contrast maps. The resulting contrast maps were then fitted to the empirical contrast maps obtained to minimise the sums of squared errors for each individual for individual-level analysis, and across individuals for the group-level analysis. Of note, rather than fitting the model to the across-participant averaged contrast map in the group-level analysis, the model minimised all the individual maps at once, allowing for a single model fit for the group without averaging away individual map features. The optimisation process was done using the *fminsearch* function of the *Optimization* toolbox in Matlab. The free parameter search was initialised with *β*_0_ = 0 and *γ*_0_ = 0. Model comparisons were performed by finding the model with lowest BIC, defined as *BIC* = *n* log(*RSS/n*) + *k* log *n* with *n* = 100^2^ = 10000 the number of timepoint per participant map, *k* the number of parameters in the model considered and *RSS* the model’s residual sum of squares.

## 5 Acknowledgments

We would like to thank John-Stuart Brittain for suggestions and comments on the analyses and R. Chris Miall for helpful comments on the manuscript. This work was supported by the European Research Council grant MotMotLearn 637488.

---

**Figure 3–Figure supplement 1.**
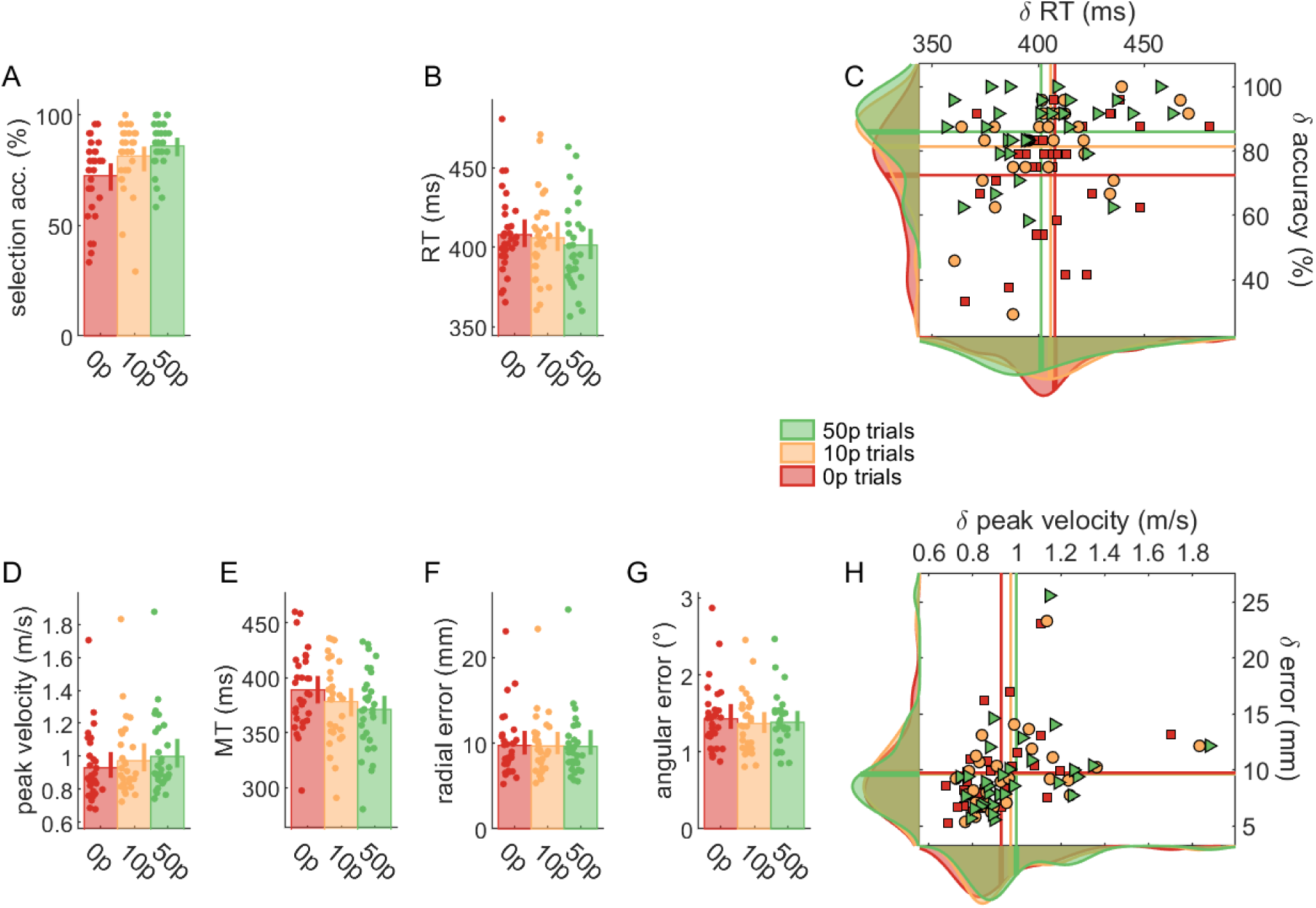
Non-normalised data for all variables in the reward-magnitude experiment.

**Figure 5–Figure supplement 1.**
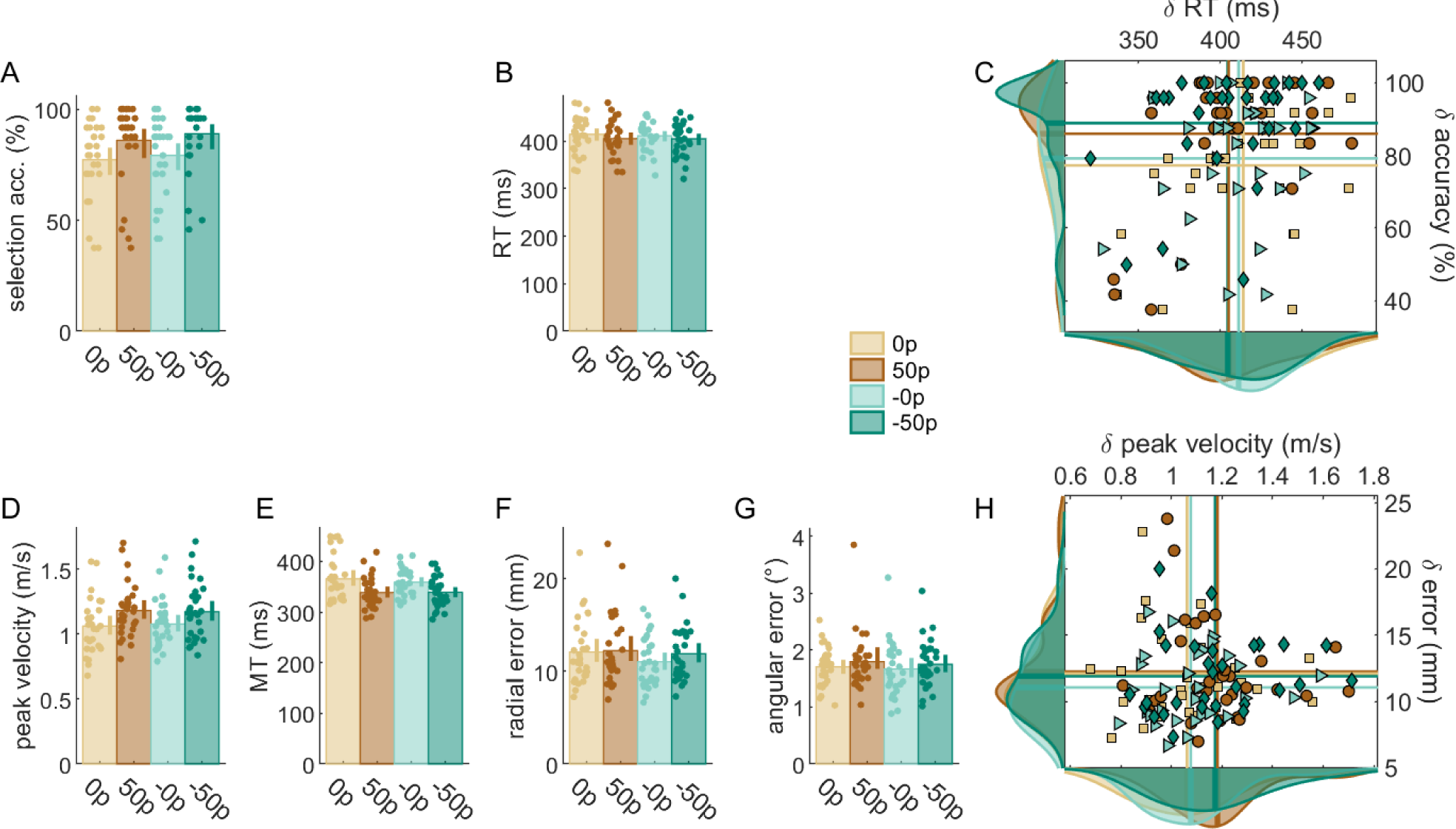
Non-normalised data for all variables in the reward-punishment experiment.

**Figure 5–Figure supplement 2.**
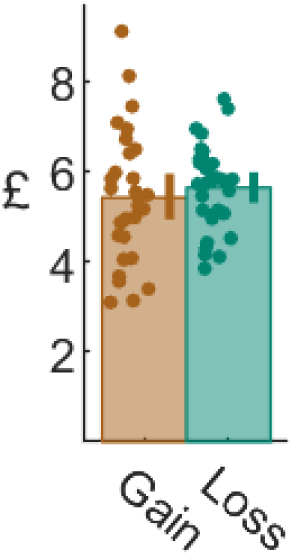
Amount of monetary gains and losses in the reward-punishment experiment. Participants earned on average the same amount of money in the rewarded block as they lost during the punishment block (see section Experimental design for details).

**Figure 7–Figure supplement 1.**
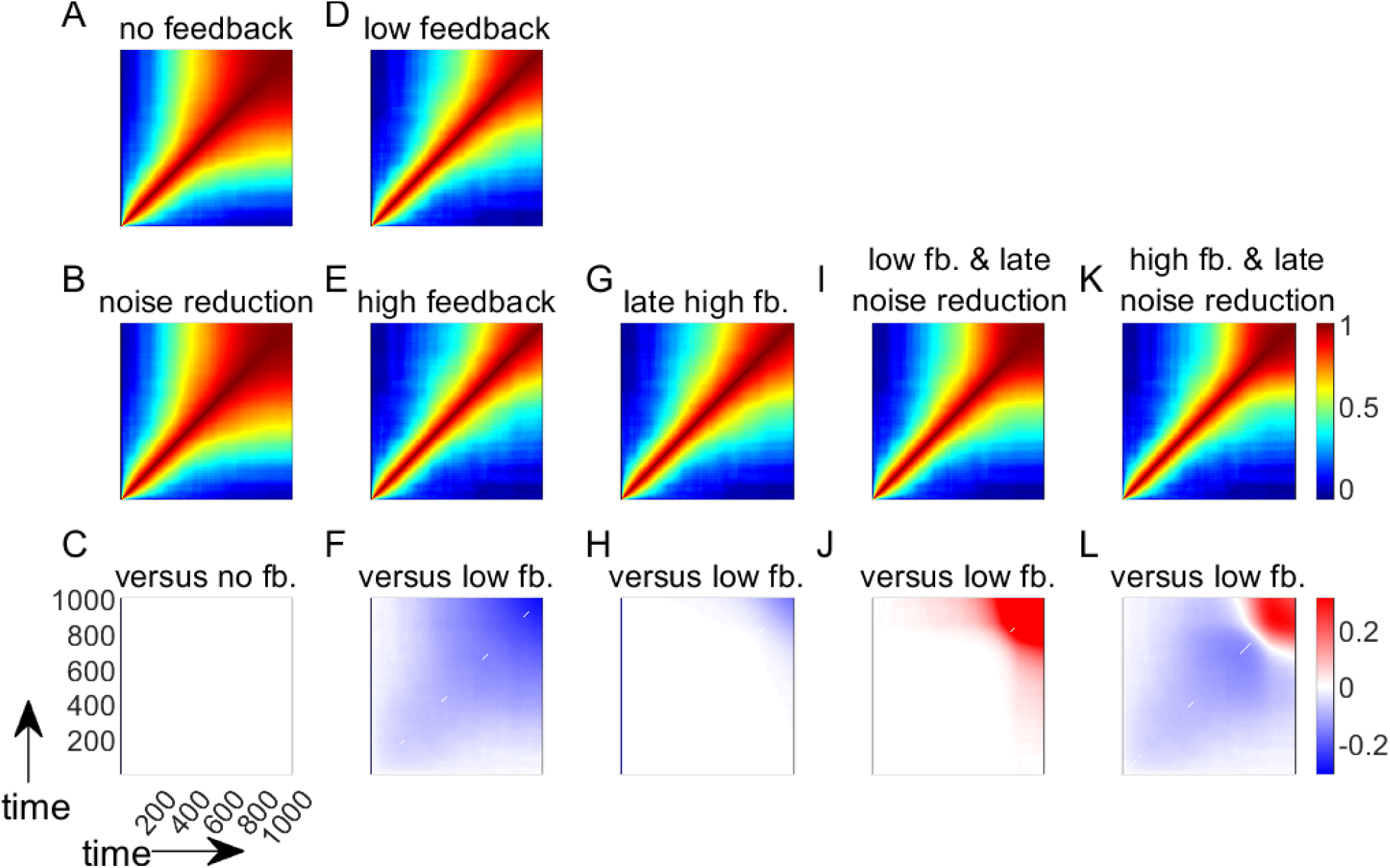
Simulations with a bell-shaped noise term to introduce signal-dependent noise.

**Figure 7–Figure supplement 2.**
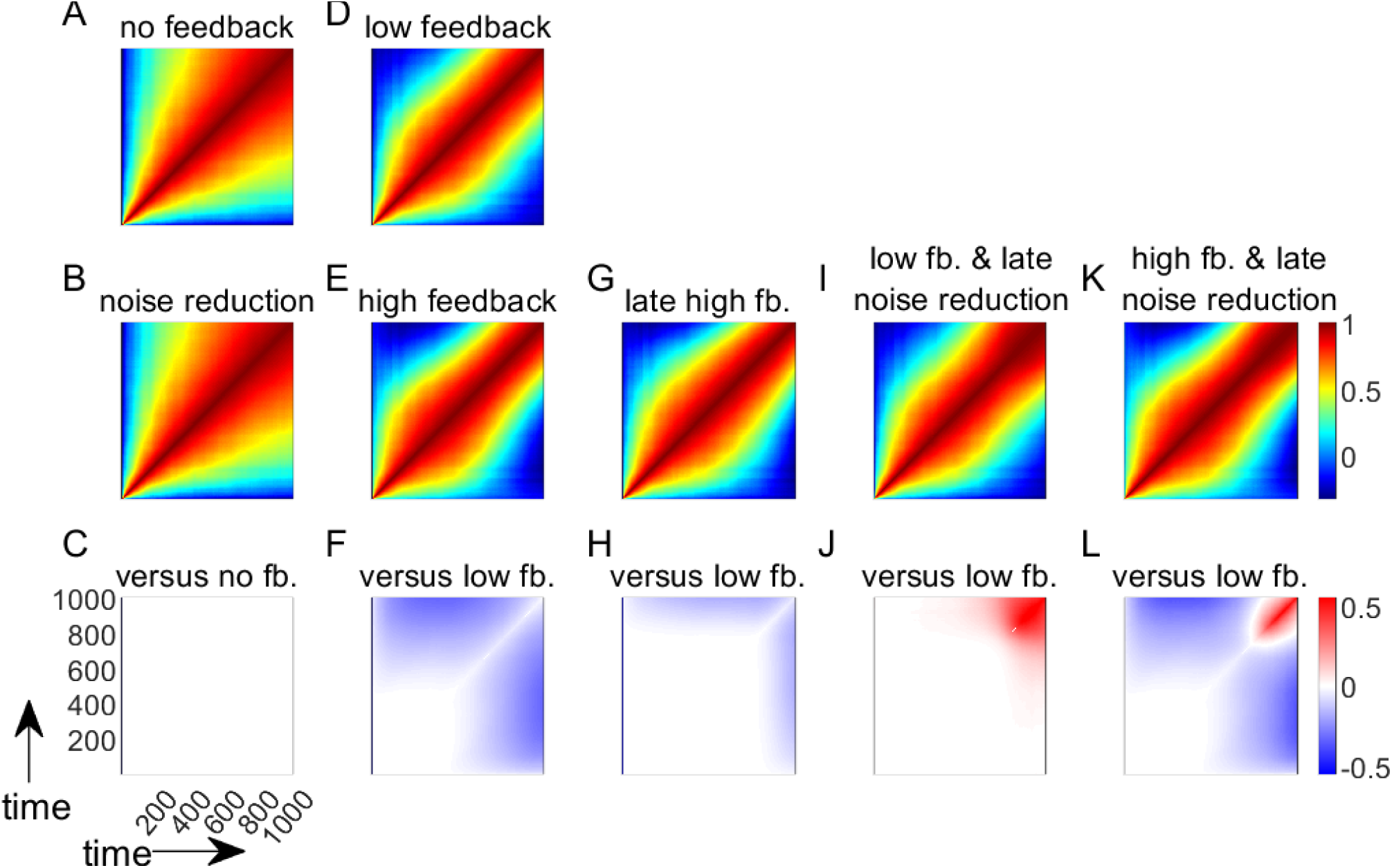
Simulations with feedback delay of 400 timesteps.

**Figure 10–Figure supplement 1.**
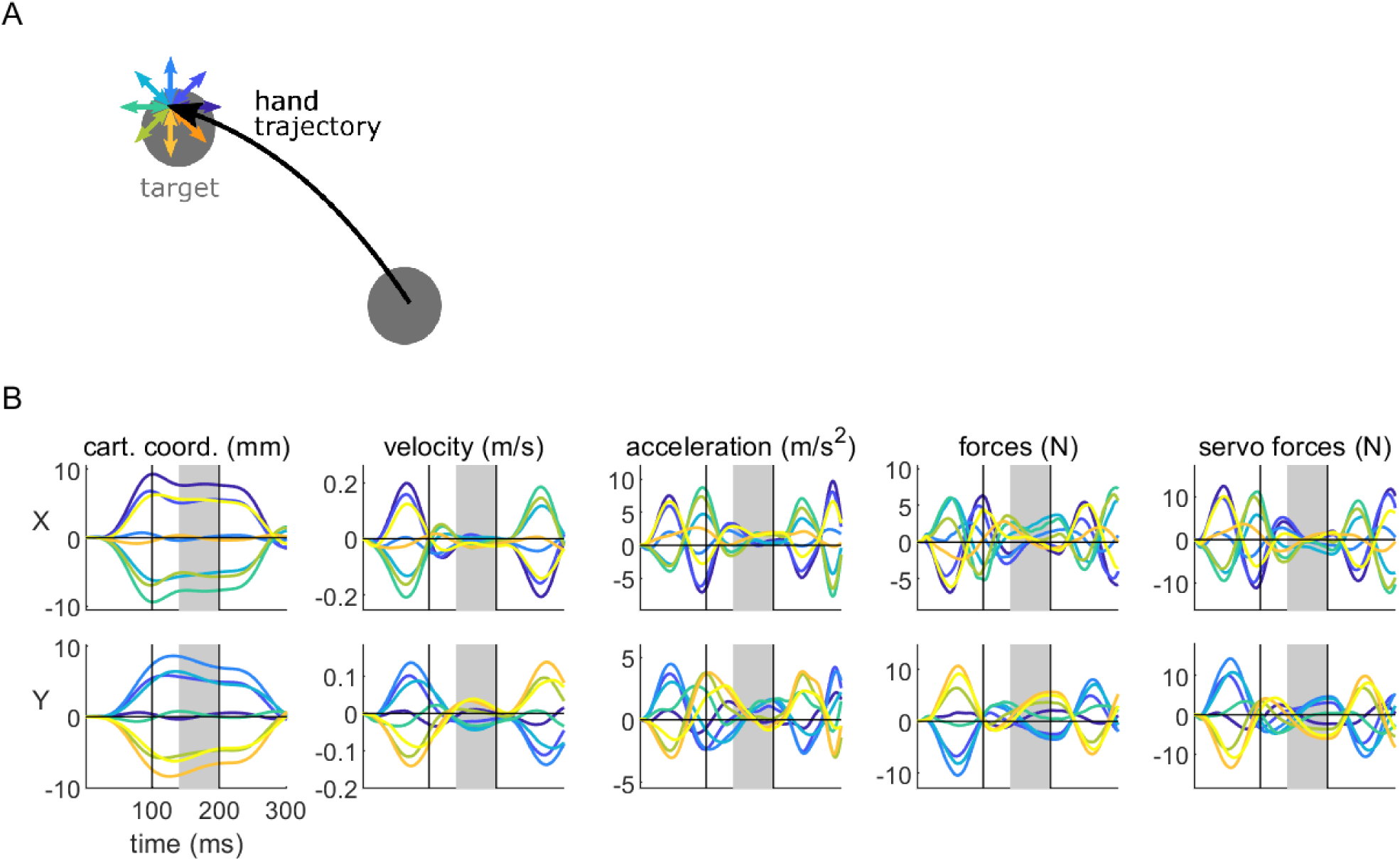
Displacement profile at the end of the reaching movement. A. Schematic of the displacement. At the end of the movement, when velocity decreased behind a threshold of 0.3 m/s, a displacement occasionally occurred in one of 8 possible directions. Each direction is represented by a colour. B. Average displacement profile over time for the first participant on the right-hand side target. The upper and lower rows represent variables in the *x* and *y* dimension, respectively. The two vertical black solid lines demark the limit between the ramp-up and plateau, and plateau and ramp-down phase. Values for each variable were taken as the average over time during the 140-200ms window (grey area), where the displacement is clamped and most stable.

**Figure 10–Figure supplement 2.**
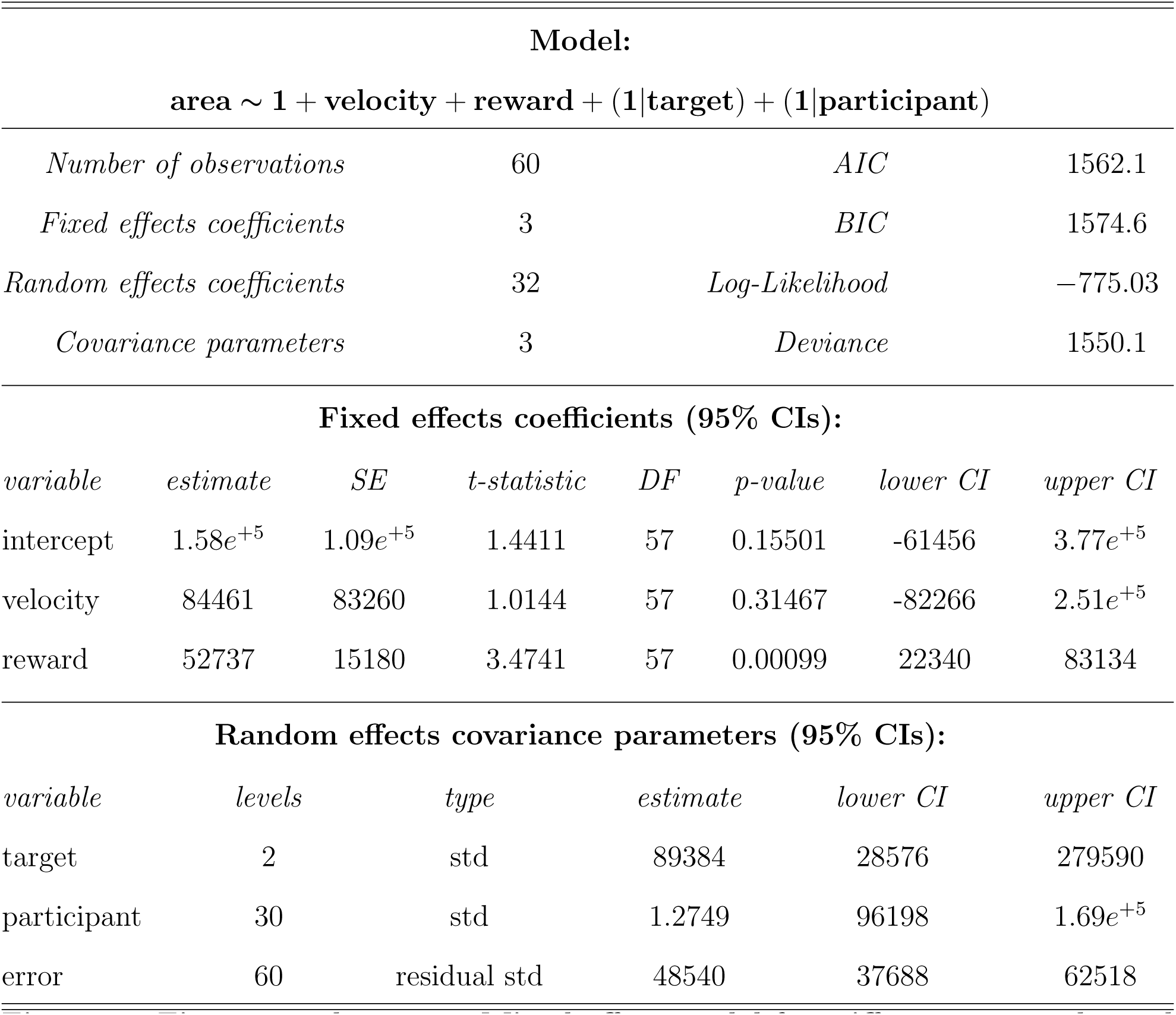
Mixed-effect model for stiffness area at the end of the reaching movement.

**Figure 10–Figure supplement 3.**
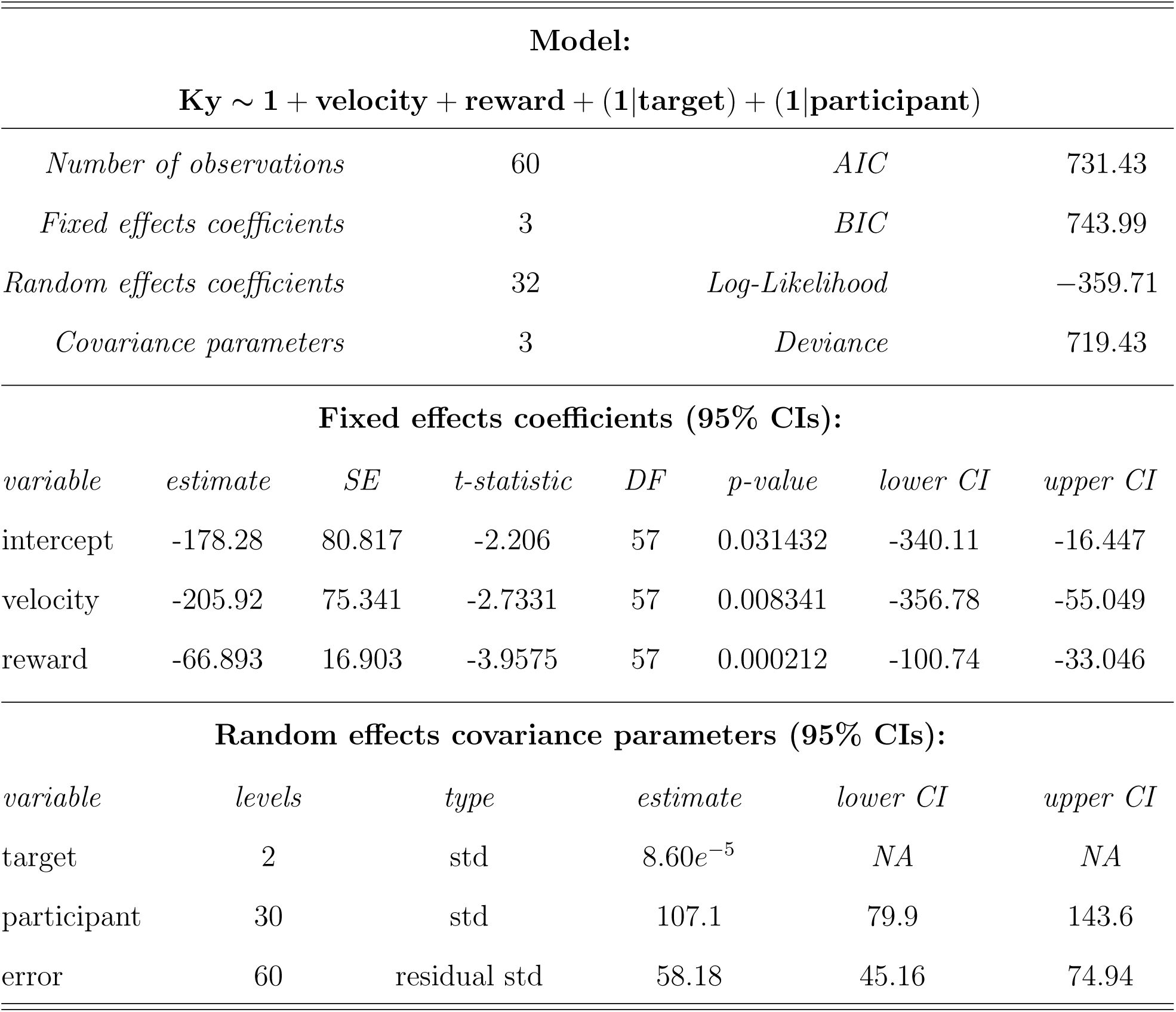
Mixed-effect model for stiffness *K*_*y*_ component at the end of the reaching movement.

**Figure 11–Figure supplement 1.**
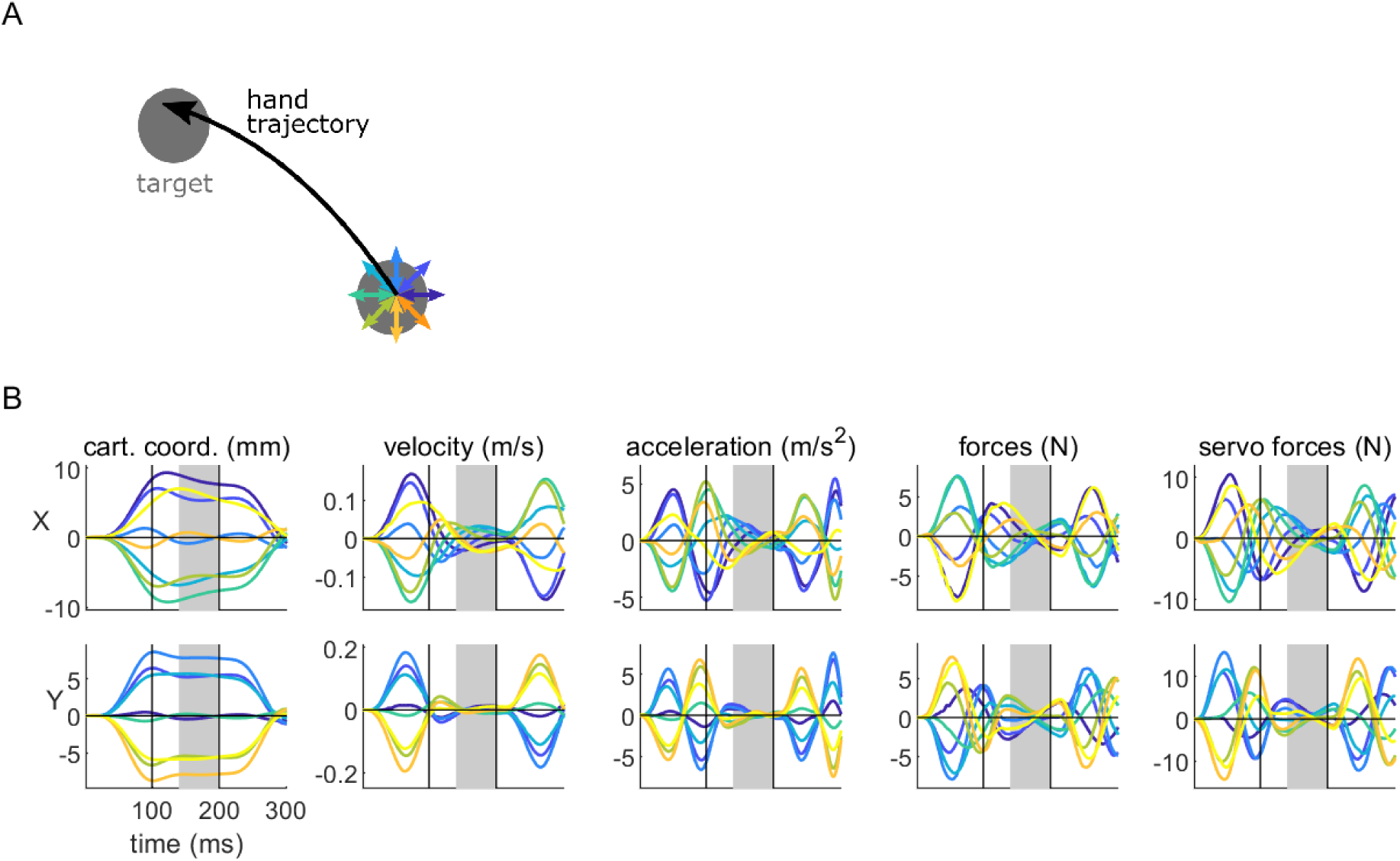
Displacement profile at the start of the reaching movement. A. Schematic of the displacement. At the start of the movement, a displacement occasionally occurred in one of 8 possible directions. Each direction is represented by a colour. B. Average displacement profile over time for the first participant. The upper and lower rows represent variables in the *x* and *y* dimension, respectively. The two vertical black solid lines demark the limit between the ramp-up and plateau, and plateau and ramp-down phase. Values for each variable were taken as the average over time during the 140-200ms window (grey area), where the displacement is clamped and most stable.

**Figure 11–Figure supplement 2.**
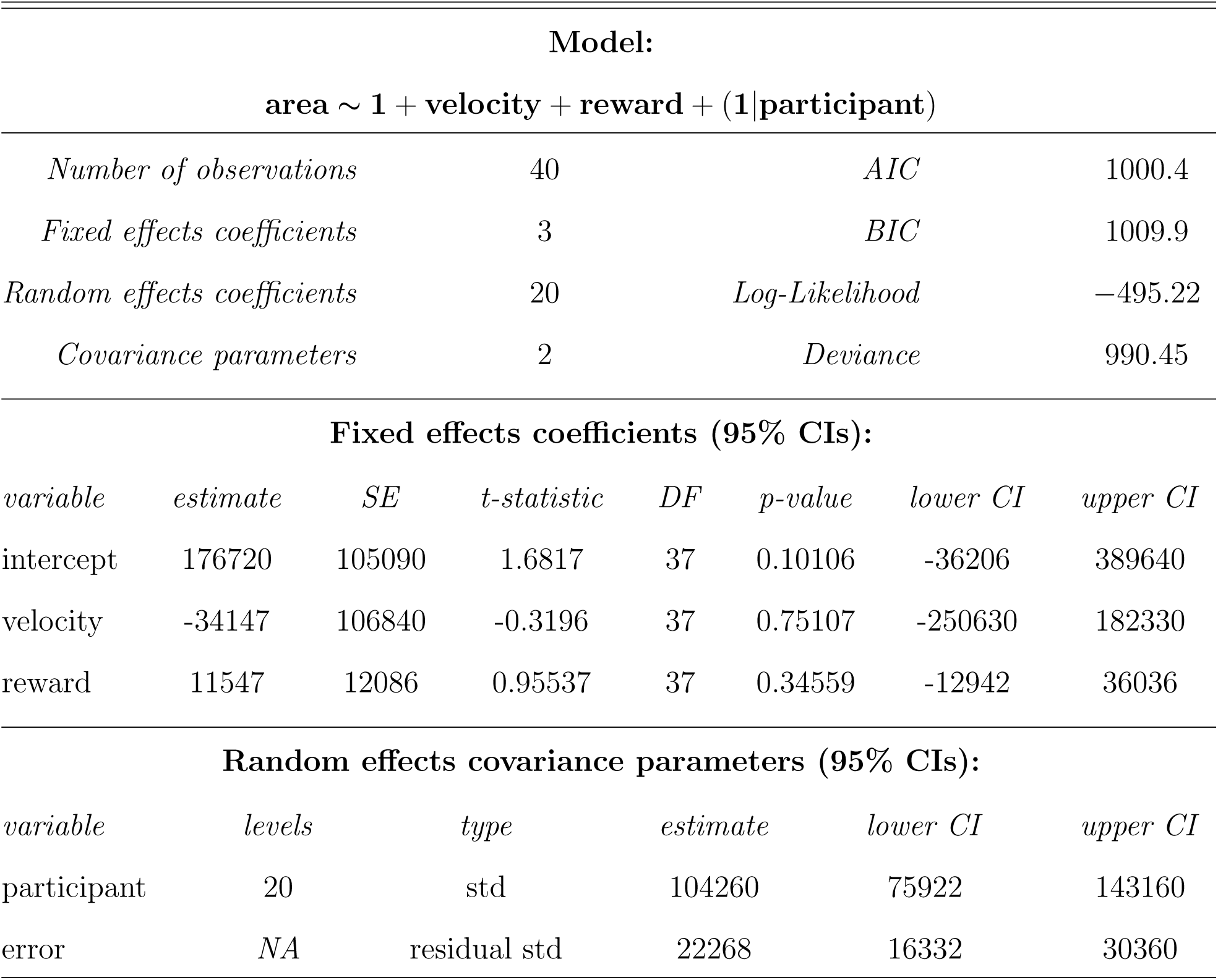
Mixed-effect model for stiffness area at the start of the movement.

**Figure 11–Figure supplement 3.**
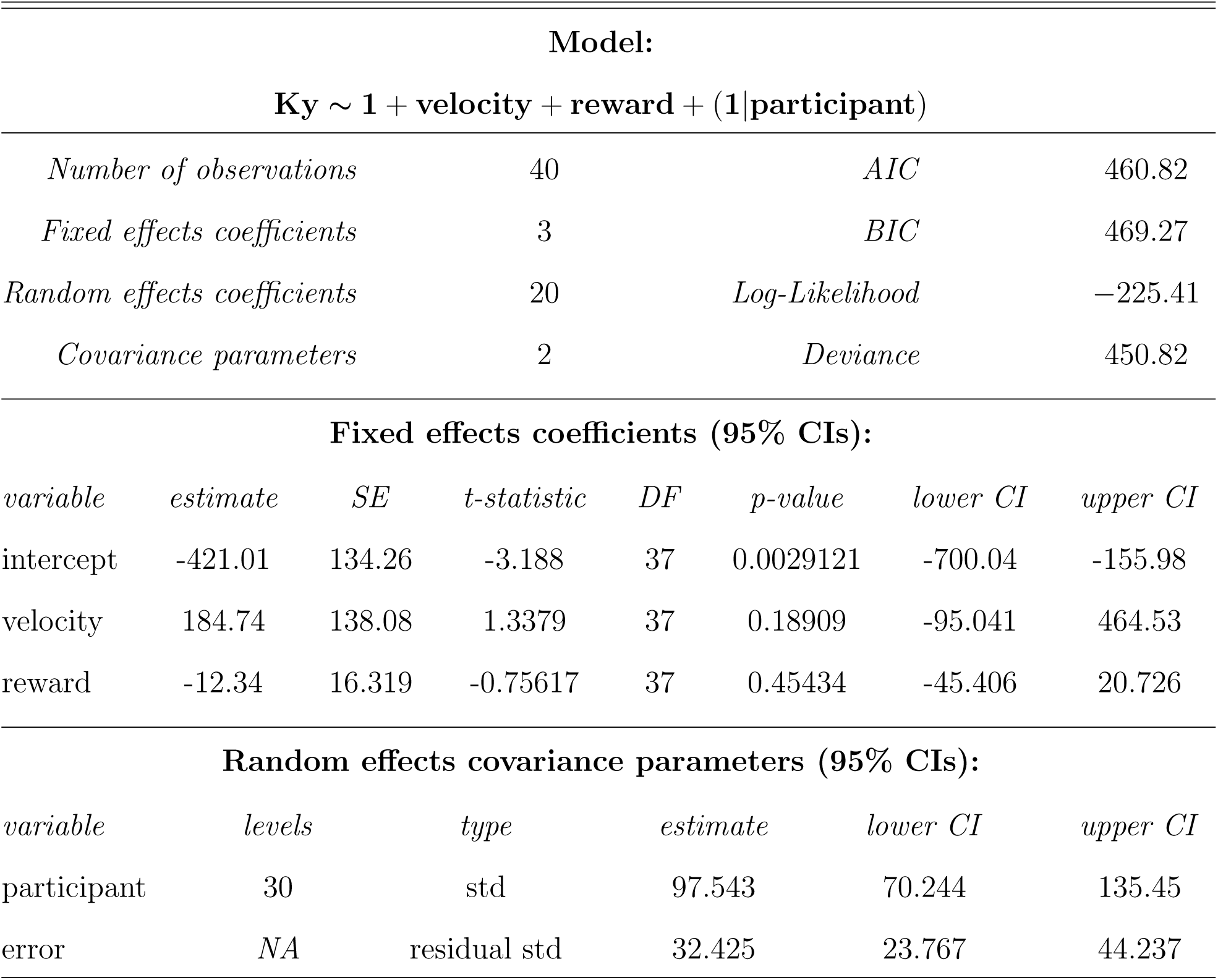
Mixed-effect model for stiffness *K*_*y*_ component at the start of the movement.

